# Lysyl oxidase promotes actin-dependent neutrophil activation and cytotoxicity in diabetes: Implications for diabetic retinopathy

**DOI:** 10.1101/2025.05.02.651525

**Authors:** Mahesh Agarwal, Sathishkumar Chandrakumar, Irene Santiago Tierno, Emma M. Lessieur, Zak R. Bollinger, Timothy S. Kern, Kaustabh Ghosh

**Affiliations:** Department of Ophthalmology, University of California, Los Angeles, CA, USA 90095; Doheny Eye Institute, Pasadena, CA, USA 91103; Molecular, Cellular, and Integrative Physiology Interdepartmental PhD Program, University of California, Los Angeles, CA, USA 90095; Department of Ophthalmology, Center for Translational Vision Research, University of California, Irvine, CA 92697; Gavin Herbert Eye Institute, University of California, Irvine, CA, USA 92697

**Keywords:** Diabetes, Lysyl oxidase, Leukocytes, Neutrophil elastase, Retinal endothelial cells, Actin

## Abstract

Activated neutrophils contribute to retinal endothelial cell (EC) death and capillary degeneration associated with early diabetic retinopathy (DR). However, the factors and mechanisms driving neutrophil activation and cytotoxicity in diabetes remain insufficiently understood. Here we show that lysyl oxidase (LOX), a collagen crosslinking and matrix stiffening enzyme that increases retinal EC susceptibility to activated neutrophils, simultaneously activates neutrophils in its alternate soluble form. Specifically, treatment of diabetic mice with LOX inhibitor β-aminopropionitrile (BAPN) prevented the diabetes-induced increase in neutrophil activation (extracellular release of neutrophil elastase and superoxide) and cytotoxicity towards co-cultured mouse retinal ECs. Mouse neutrophils and differentiated (neutrophil-like) human HL-60 cells treated with recombinant LOX alone exhibited similar activation and cytotoxicity. Mechanistically, this LOX-induced neutrophil activation was associated with biphasic F-actin remodeling, with the initial and rapid (<15 min) F-actin depolymerization followed by a significant increase in F-actin polymerization and polarization. Preventing the initial F-actin depolymerization blocked LOX-induced neutrophil activation and cytotoxicity towards co-cultured retinal ECs. Finally, we show that this biphasic F-actin remodeling is essential for LOX-induced membrane clustering of azurophilic granule marker CD63 and NADPH organizer p47 that are associated with extracellular release of neutrophil elastase and superoxide, respectively. By revealing a causal and previously unrecognized link between LOX and actin-dependent neutrophil activation in diabetes, these findings provide fresh mechanistic insights into the proinflammatory role of LOX in early DR that goes beyond its canonical matrix-stiffening effects.

## INTRODUCTION

Inflammation plays a causal role in retinal endothelial cell (EC) dysfunction associated with diabetic retinopathy (DR), a major microvascular and vision-threatening complication of diabetes (1, 2). Among the key proinflammatory mediators of retinal EC dysfunction are activated neutrophils that bind to ECs and release abundant neutrophil elastase (NE) and reactive oxygen species (ROS) extracellularly (3–5). While these neutrophil-derived factors are primarily aimed at fighting pathogens, their high extracellular levels cause retinal EC apoptosis (3, 4). Progressive loss of retinal ECs eventually leads to retinal capillary degeneration, an important clinically recognized lesion of early DR. Thus, identifying the proinflammatory factors and intracellular mechanisms that drive neutrophil activation has important therapeutic implications for early management of DR.

Recent work has identified lysyl oxidase (LOX) as a potent proinflammatory factor that promotes retinal EC dysfunction in early DR (6–8). LOX is a copper-dependent enzyme known primarily for its ability to crosslink collagen/elastin in extracellular matrix, thereby stiffening it (9, 10). Consistent with this mechanical role of matrix-localized LOX, we recently showed that diabetes-induced LOX upregulation in retinal ECs promotes retinal capillary stiffening and activation of mechanosensitive NF-κB in retinal ECs (6–8). This, in turn, increases endothelial ICAM-1 expression, neutrophil-EC adhesion, and increased EC susceptibility to neutrophil-/NE-induced cytotoxicity (6–8). These findings align with the known proinflammatory role of matrix LOX in other vascular complications such as pulmonary edema and atherosclerosis (11–13).

However, accumulating evidence points to an important, albeit less recognized, role of the alternate soluble form of LOX and its family members LOX-like proteins 1-4 (LOXL1-4). These observations, made primarily in cancer cells, implicate soluble LOX in the transcriptional, epigenetic, and cytoskeletal regulation of epithelial-to-mesenchymal transition (EMT), a crucial process that promotes tumor progression and metastasis (9, 14). The intracellular effects of soluble LOX on other cells and pathologies are, however, poorly understood. Since retinal endothelial LOX expression increases in diabetes and pharmacological LOX inhibition reduces leukocyte-EC adhesion and retinal EC death (7), we asked whether, and if so how, soluble LOX produces intracellular changes in neutrophils that facilitate their activation and cytotoxicity towards retinal ECs.

Among the intracellular mechanisms implicated in neutrophil activation (NE and superoxide release) by proinflammatory factors is actin remodeling (15–17). Past studies looking at the role and regulation of neutrophil actin organization have, however, produced varying results. For instance, Kutsuna et al. showed that TNF-treated human neutrophils undergo rapid (within 30 min) F-actin depolymerization in vitro (18) while another study reported an opposite trend (19). Further, while increasing F-actin polymerization in human neutrophils was found to inhibit their activation (CD11b expression) in diabetes (20), decreasing F-actin polymerization (using Arp2/3 inhibitor) in mouse neutrophils reduced sepsis-induced neutrophil extracellular trap (NET) formation and associated lung damage (21). Although these findings implicate F-actin remodeling as an important mediator of neutrophil activation and function, its ability to exert differential (stimulatory vs inhibitory) effects underscores the need to systematically investigate the cytoskeletal regulation of neutrophil activation by proinflammatory factors such as LOX.

Using a mouse model of DR, which exhibits both retinal LOX upregulation and neutrophil activation, and complementary cultures of isolated mouse neutrophils and differentiated (neutrophil-like) HL-60 cells, we show that LOX induces neutrophil activation that is marked by increased NE and superoxide release, greater cytotoxicity towards retinal ECs, membrane aggregation of CD63-rich azurophilic granules and NADPH organizer p47^phox^, and biphasic F-actin dynamics affecting both F-actin polymerization and polarization. Importantly, our studies reveal that dynamic F-actin remodeling is both necessary and sufficient to induce neutrophil activation and cytotoxicity.

## RESEARCH DESIGN AND METHODS

### Experimental Animals

All animal procedures were performed in accordance with the Association for Research in Vision and Ophthalmology (ARVO) Statement for the Use of Animals in Ophthalmic and Vision Research and approved by University of California Los Angeles Institutional Animal Care and Use Committee. To induce diabetes in adult (8-week-old) male C57BL/6J mice (Jackson Laboratory, Bar Harbor, ME, USA), i.p. injections of freshly prepared streptozotocin solution (STZ; MP Biomedicals, Irvine, CA; 60 mg/kg body weight in 10 mM citrate buffer; pH 4.5) were administered daily for five consecutive days. Mice were classified as diabetic if their fasting blood glucose was >275 mg/dL two weeks after the last STZ injection, with this time point being considered as the onset of overt diabetes. Age-matched normal C57BL/6J mice that only received citrate buffer served as non-diabetic controls. Some diabetic mice were treated with a specific and irreversible LOX inhibitor β-aminopropionitrile (22) (BAPN; 3 mg/kg through drinking water; Catalog no. A3134-5G; Sigma-Aldrich, St. Louis, MO, USA) for 10 weeks (from the onset of overt diabetes) prior to euthanasia and collection of eyes, femur, and tibia for further analyses. Nondiabetic, diabetic, and diabetic mice receiving BAPN are henceforth referred to as ND, D, and D+BAPN mice.

### Neutrophil Isolation

Neutrophils were isolated from the bone marrow of femur and tibia of euthanized ND, D, or D+BAPN mice using the EasySep Mouse Neutrophil Enrichment Kit (Stemcell Technologies, Vancouver, Canada, Catalog no. 19762), as per manufacturer’s protocol that results in >90% pure neutrophils.

### Cell Culture and Treatments

Human promyelocytic leukemia cell line (HL-60) was purchased from ATCC (Manassas, VA, USA) and cultured in RPMI 1640 medium (Gibco™-Thermo Fisher Scientific) supplemented with 10% fetal bovine serum (FBS; Catalog no. SH30396.03, Hyclone, Logan, UT) and Penicillin-Streptomycin (Gibco™-Thermo Fisher Scientific). HL-60 cells were differentiated with 1.4% (v/v) DMSO for 5 days to acquire the functional properties of neutrophils (23). The differentiated HL-60 (dHL-60) cells, widely used as an alternative to primary neutrophils (24–26), or the isolated mouse bone marrow neutrophils were treated with recombinant human LOX (Catalog no. 228-20774-2, Ray Biotech, USA) or recombinant mouse LOX (Catalog no. LS-G 15002-50, LS bio, MA, USA), respectively, at the indicates doses or for the indicated durations prior to use in subsequent neutrophil cytotoxicity or activation assay. For studies looking at the role of actin remodeling, dHL-60 cells were also treated with (a) LOX + actin stabilizing drug Jasplakinolide (27) (1 µM; Catalog no. 420107 Sigma-Aldrich, St. Louis, MO, USA), or (b) actin depolymerizing drug cytochalasin D (cytoD) alone (2.5 µM; Catalog no. 2502551, MilliporeSigma St. Louis, MO, USA). Cells were treated for 6h prior to use in subsequent neutrophil activation or cytotoxicity assay. Human retinal endothelial cells (HRECs) were purchased from Cell Systems Corp. (Catalog no. ACBRI 181, Kirkland, WA, USA;) and cultured as previously reported (8). Mouse retinal endothelial cells (mRECs) were purchased from Cell Biologics Inc. (Catalog no. C57-6065, Chicago, IL, USA;) and cultured in vendor recommended medium (Catalog. no. M1168; Cell Biologics). HRECs and mRECs were used until passage 10.

### Neutrophil-/dHL-60-mediated Cytotoxicity Toward Retinal Endothelial Cells

Neutrophil-/dHL-60-mediated cytotoxicity toward RECs was assayed as previously described (3). Briefly, three sets of co-cultures were performed: 1) bone marrow neutrophils from ND, D, or D+BAPN mice (n=6/group) co-cultured with mRECs, 2) bone marrow neutrophils from ND mice treated with recombinant mouse LOX (75 ng/mL; Catalog no. LS-G 15002-50, LS bio, MA, USA) for 6h prior to co-culture with mRECs, and 3) dHL-60 cells treated with recombinant human LOX±Jasplakinolide (1µM) or cytoD alone (2.5 µM) for 6h prior to co-culture with HRECs. All neutrophil/dHL-60 co-cultures with RECs were maintained at a cell ratio of 1:3 for 16 h. In some studies, neutrophil elastase inhibitor Sivelestat (50 µM) (3, 28) (Catalog no. ab146184 Abcam, Cambridge, MA, USA) was added to HRECs 30 min prior to addition of dHL-60 and maintained for the entire duration (16 h) of co-culture. All experimental conditions were maintained in triplicates. Next, the co-culture was detached using trypsin and labeled with anti-human CD144 (Catalog no. BD541567, BD biosciences San Deigo, USA) or anti-mouse CD144 (Catalog no. 138005 Biolegends, San Diego, USA) to identify RECs, together with Annexin V (Catalog no. BD 550911 BD biosciences San Deigo, USA) and Propidium Iodide ( Catalog no. 537059 Sigma-Aldrich, St. Louis, MO, USA) to assess REC death by flow cytometry. A total of 10,000 events were counted for each sample. Results were analyzed by Flow Jo 10.7.2.

### Western Blot

To determine the protein expression levels of NE and CD11b, equal amounts (20 or 40 µg) of total protein were separated in 4-15% Mini-PROTEAN® TGX precast protein gels (Biorad, Hercules, CA, USA) and transferred to nitrocellulose membrane before probing with primary anti-NE (catalog no. NB300-605; Novus Biologicals, San Diego, CA, USA), anti-CD11b (Catalog no. NB110-89474, Novus Biologicals, San Diego, CA, USA,), and anti-GAPDH (loading control; Catalog no. G9545; Sigma-Aldrich), followed by appropriate secondary antibody conjugated to horseradish peroxidase (Catalog no. PI-1000 and PI-2000; Vector Laboratories). Protein bands were detected using a SuperSignal™ West Dura Extended Duration Substrate (Catalog no. 34075; Thermo Fisher Scientific, Waltham, MA, USA) and imaged using ChemiDoc XRS+ System (Biorad). Targets were assessed from technical duplicates obtained from three independent experiments. Densitometric analysis was performed using ImageJ.

### Extracellular NE Activity

Activity of extracellular NE released by mouse neutrophils or dHL-60 cells was measured using the EnzChek Elastase Assay Kit protocol (Catalog no. E12056, Invitrogen) (3). Briefly, equal number of untreated or treated mouse neutrophils or dHL-60 cells suspended in Ca^2+^ buffer (136 mM NaCl; 4.7 mM KCl; 1.2 mM MgSO_4_; 1.1 mM CaCl_2_; 1.2 mM KH_2_PO_4_; 5 mM NaHCO_3_; 5.5 mM Glucose; 20 mM HEPES in ddH_2_O) were briefly stimulated with fMLP (10 nM; 5 min) prior to centrifugation. Equal volume of the (cell-free) supernatant was then added to a 96-well plate pre-loaded with elastase substrate (quadruplicates/condition) and incubated for 30 min prior to fluorescence measurement (505/525 nm) using a spectrometer (Molecular device Spectromax iD5).

### Superoxide Generation

Superoxide generation by mouse neutrophils or dHL-60 cells was measured using DHE dye (Catalog no. D11347, Invitrogen) (29). Briefly, equal number of untreated or treated mouse neutrophils or dHL-60 cells were suspended in Ca^2+^ buffer (136 mM NaCl; 4.7 mM KCl; 1.2 mM MgSO_4_; 1.1 mM CaCl_2_; 1.2 mM KH_2_PO_4_; 5 mM NaHCO_3_; 5.5 mM Glucose; 20 mM HEPES in ddH_2_O) containing 10 µM DHE dye for 30 min prior to brief stimulation with fMLP (10 nM; 5 min). Next, equal number of cells were added to a 96-well plate (quadruplicates/condition) for fluorescence measurement (518/606 nm) using a spectrometer (Molecular device Spectromax iD5).

### RT-qPCR

Total RNA was isolated from mouse retina (n=5/group) or dHL-60 cells (biological triplicates/condition) using Direct-zol RNA Miniprep kit (Zymo Research, Irvine, CA, USA). Next, equal amount (1 µg) of total RNA was converted to cDNA using a High-Capacity RNA-to-cDNA kit (catalog no. 4387406; Thermo Fisher Scientific-Applied Biosystems™) before amplification with gene/species-specific TaqMan primers for LOX (Mm00495386_m1; Hs00942480_m1) in QuantStudio™ 5 Real-Time PCR system. Target gene expression was normalized to the house keeping gene GAPDH (Mm99999915_g1; Hs02786624_g1) and relative expression was determined using the comparative Delta Delta Ct (DDCt) method. Negative control was performed for each reaction assay plate, with each reaction run in technical duplicate.

### Immunofluorescence Staining for CD63 and p47^phox^

Mouse neutrophils or dHL-60 cells were fixed using a previously published protocol (30). Briefly, cells were suspended in a fixation buffer (280 mM KCl, 2 mM MgCl_2_, 4 mM EGTA, 40 mM HEPES, 0.4% bovine serum albumin/BSA, 640 mM sucrose, 7.4% (w/v) formaldehyde, pH 7.5) for 20 min at RT prior to rinsing with an intracellular buffer composed of 140 mM KCl, 1 mM MgCl_2_, 2 mM EGTA, 20 mM HEPES, 0.2% BSA, pH 7.5, as previously reported (30). The fixed cells were then blocked with 2% (w/v) BSA (in PBS) for 1h at RT prior to incubation with anti-CD63 (Catalog no. PA5-100713, Invitrogen) or anti-p47^phox^ (Catalog no. PA1-9073, Invitrogen) in 2% BSA (in PBS) for overnight at 4°C. Next, cells were rinsed twice with intracellular buffer prior to incubation with an appropriate Alexa Fluor-conjugated secondary antibody (Jackson lab) in 2% BSA (in PBS) for 1h at 4°C. Immunolabeled cells were finally rinsed twice with intracellular buffer prior to counterstaining with phalloidin to visualize F-actin, as described in the following section.

### F-actin staining

Immunolabeled mouse neutrophils or dHL-60 cells were incubated with rhodamine-conjugated phalloidin (130 nM; ThermoFisher) in the aforementioned intracellular buffer containing 0.2% (v/v) Triton X-100 for 30 min at RT. Next, labeled cells were rinsed with intracellular buffer and mounted on Vectabond-treated coverslides for imaging using Zeiss LSM 710 confocal microscope (63X/NA 1.2 objective). For the time course studies, dHL-60 cells were fixed at predetermined time points, labelled with rhodamine-phalloidin, and mounted on Vectabond-treated coverslides for imaging using Zeiss Axio Observer (63X/NA 1.4 objective) as per the aforementioned protocol.

### Aggregation/Polarization index

The aggregation of CD63-containing azurophilic granules or NADPH organizer p47^Phox^ as well as the polarization of F-actin in mouse neutrophils or dHL-60 cells were quantitatively analyzed by measuring the distance between the cell’s geometric centroid and the weighted centroid of the cell’s fluorescence signal. Note that in a cell with uniformly distributed azurophilic granules, p47^Phox^, or F-actin, the geometric and weighted centroids should colocalize. However, these centroids will separate (that is, cell’s polarization index will increase) as the cell’s granules, p47^Phox^, or F-actin preferentially aggregate/polarize at one end, leading to their non-homogenous fluorescent labeling in cells. Please refer to Fig. 2E for a schematic illustration.

### Stiffness Measurements

Mouse neutrophils or dHL-60 cells were mildly fixed with 0.5% paraformaldehyde for 10 min at room temperature prior to stiffness measurement using NanoWizard^®^ 4 XP BioScience atomic force microscope (AFM; Bruker Nanotechnologies, Santa Barbara, CA) fitted with a pre-calibrated SAA-SPH-1UM probe (spring constant 0.25 N/m) containing a 1 µm-radius hemispherical silicon nitride tip (Bruker AFM Probes, CA, USA). The AFM was coupled with a Zeiss Axiovert phase contrast microscope to facilitate sample visualization and measurement. Stiffness was measured in the contact mode force spectroscopy mode by applying a 200 pN (set point) indentation force. Force curves from multiple cells (n ≥ 20/condition) were analyzed using JPK Data Processing Software.

### Statistics

Data analysis was performed using OriginPro2022 software (OriginLab Corporation Massachusetts, USA). Statistical differences between three or more groups were assessed using a one-way analysis of variance (ANOVA) followed by Tukey’s or Dunnett’s *post hoc* multiple comparisons test, based on whether or not the data was normally distributed. Studies comparing two experimental groups were subjected to two-tailed unpaired Student’s *t* test. Results were considered significant if *p*<0.05.

## RESULTS

### LOX mediates diabetes-induced neutrophil activation and cytotoxicity towards retinal ECs

We previously showed that matrix-localized LOX promotes retinal EC apoptosis in diabetes by increasing EC susceptibility to neutrophil-induced cytotoxicity (7). To determine whether LOX, in its alternate soluble form, could directly promote neutrophil cytotoxicity, here we isolated neutrophils from nondiabetic (ND), 10-week diabetic (D), or diabetic mice treated with LOX inhibitor BAPN, which prevents diabetes-induced retinal LOX upregulation (**Fig. 1A**), and co-cultured them with mRECs for 16h. Flow cytometry-based analysis of Annexin V-labeled mRECs **(Supplementary** Fig. 1) revealed that the significant increase in mREC apoptosis caused by neutrophils from diabetic mice is blocked by BAPN treatment (**Fig. 1B**). This ability of LOX inhibitor BAPN to suppress diabetes-induced neutrophil cytotoxicity towards retinal ECs was consistent with the loss of neutrophil activation, as judged by prevention of increase in neutrophil elastase (NE) expression (**Fig. 1C**), extracellular NE activity (**Fig. 1D**), CD11b surface expression **(Supplementary** Fig. 2A), and superoxide generation **(Supplementary** Fig. 2B). Notably, the increase in neutrophil activation or cytotoxicity towards mRECs was not seen at the earlier four weeks’ duration of diabetes (**Fig. 1E, F**) when retinal LOX levels did not increase (**Fig. 1G**). Together, these findings raise the possibility that LOX upregulation in the retina plays a causal role in diabetes-induced neutrophil activation and cytotoxicity towards retinal ECs. The importance of retinal LOX in neutrophil activation is further supported by our observation that LOX expression in neutrophils remains unaltered in diabetes **(Supplementary** Fig. 2C).

**Figure 1:**
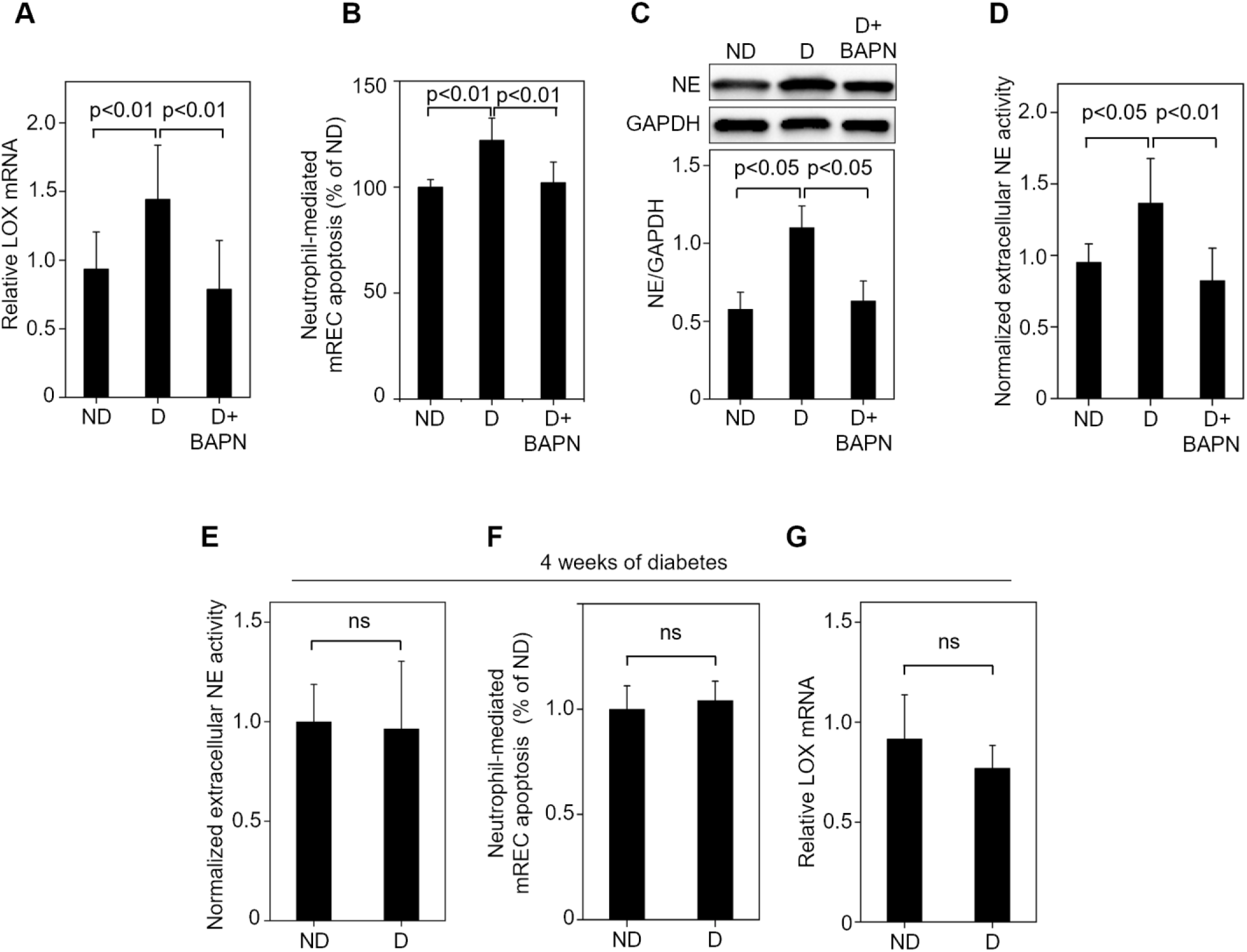
LOX mediates diabetes-induced neutrophil activation and cytotoxicity towards retinal ECs. **(A)** RT-qPCR analysis of mouse whole retina indicates that LOX inhibitor BAPN prevents the (n=6 mice/group; 10 wk diabetes) diabetes-induced increase (by 1.5-fold; p<0.01) in LOX mRNA level. **(B)** Flow cytometry-based analysis of FITC-Annexin V-labeled mRECs following 16h co-culture with neutrophils isolated from nondiabetic (ND) mice or diabetic (D) mice ± LOX inhibitor BAPN (D+BAPN; 3 mg/kg BW) (n=6 mice/group; 10 wk diabetes) revealed that the significant increase in mREC apoptosis caused by neutrophils from D mice is prevented by LOX inhibition. **(C)** Representative Western blot bands and cumulative densitometric analysis of mouse neutrophils (n=6 mice/group; 10 wk diabetes) indicate that the diabetes-induced ∼2-fold increase (p<0.05) in neutrophil elastase (NE) protein expression is prevented by LOX inhibition (using BAPN). **(D)** fMLP (10 nM) stimulation of neutrophils isolated from ND, D, or D+BAPN mice (n=6 mice/group; 10 wk diabetes) and subsequent EnzChek^™^ elastase assay revealed that the diabetes-induced ∼40% increase (p<0.05) in extracellular NE activity is prevented by LOX inhibition. **(E)** fMLP (10 nM) stimulation of neutrophils isolated from ND or 4-wk D (four weeks’ duration of diabetes) mice (n=7 mice/group) and subsequent EnzChek^™^ elastase assay indicate a lack of change in extracellular NE activity in 4-wk D mice. **(F)** Flow cytometry-based analysis of FITC-Annexin V-labeled mRECs following co-culture with mouse neutrophils isolated from ND or 4-wk D mice (n=7 mice/group) revealed no increase in neutrophil-mediated mREC apoptosis in shorter-term diabetes. **(G)** RT-qPCR analysis of mouse whole retina indicates that LOX mRNA levels do not increase in 4-wk D mice (n=7 mice/group). Bar graphs indicate mean ± SD.

### LOX induces neutrophil activation and CD63 aggregation

To determine whether LOX alone is sufficient to induce neutrophil activation, neutrophils isolated from ND mice were treated with varying doses of recombinant LOX for 6h. Our findings revealed that 75 ng/mL LOX maximally increases extracellular NE activity (by 1.4 fold; p<0.05) (**Fig. 2A, Supplementary** Fig. 3A). Further, although LOX induced neutrophil activation earlier at 30 min, the increase in extracellular NE activity (by 1.5 fold; p<0.05) and superoxide generation (by ∼1.25 fold; p<0.05) became significant later at 6h (**Supplementary** Figs. 3B, C). Predictably, this LOX-induced neutrophil activation was associated with a significant increase (by 1.4 fold, p<0.05) in cytotoxicity towards mRECs (**Fig. 2B**).

**Figure 2.**
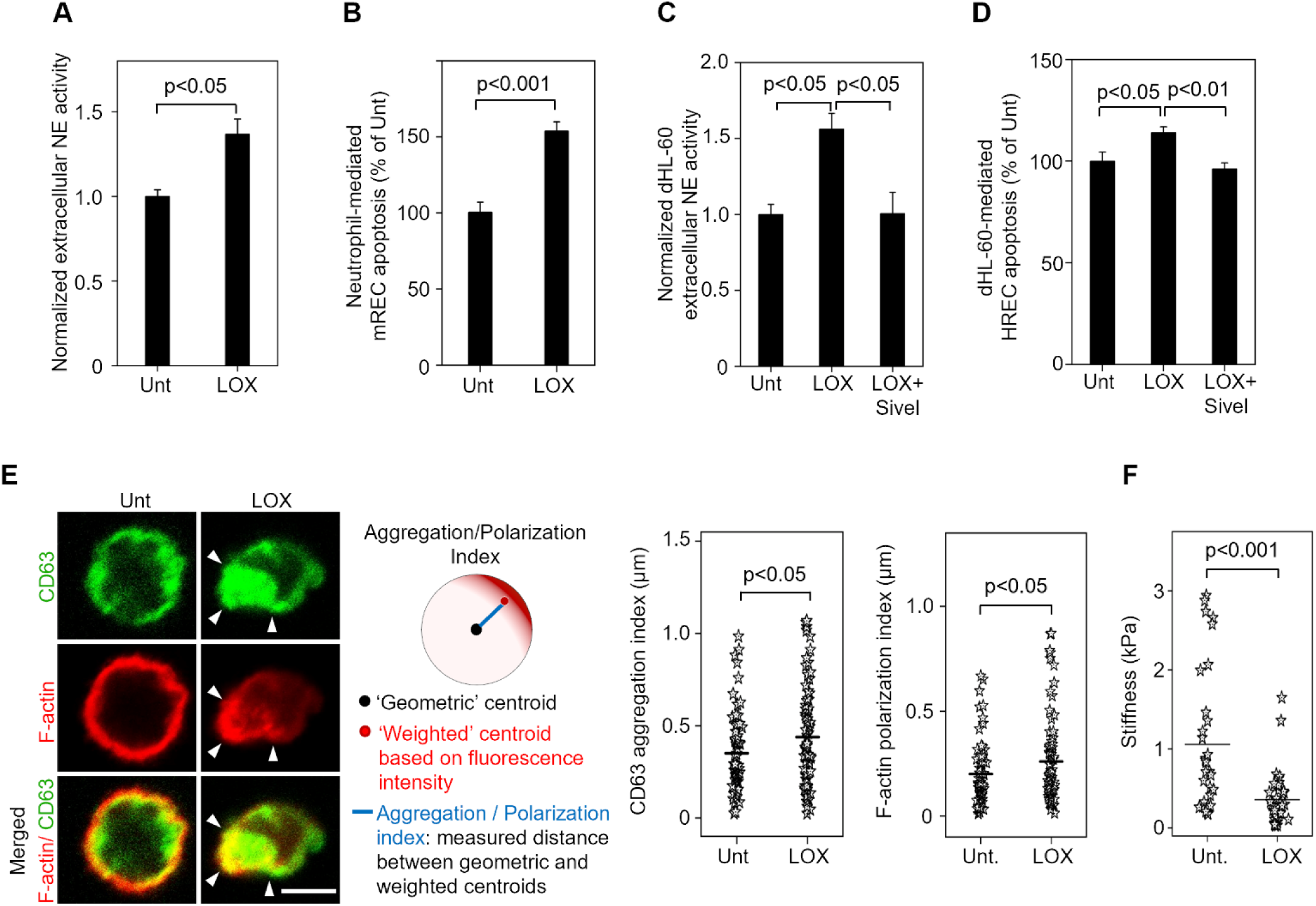
LOX induces neutrophil activation and CD63 aggregation. **(A)** Neutrophils isolated from ND mice were either left untreated (Unt.) or treated with 75 ng/mL (6h) recombinant mouse LOX before brief stimulation with fMLP (10 nM) and supernatant collection. Fluorometric analysis of the supernatant/EnzChek^™^ elastase substrate mix revealed a 40% increase (p<0.05) in extracellular NE activity in LOX-treated neutrophils. **(B)** Flow cytometry quantification of the total number of FITC Annexin V-labeled mRECs following 16h of co-culture with LOX-treated (75 ng/mL; 6h) ND neutrophils revealed a 1.5-fold increase (p<0.001) in mREC apoptosis caused by LOX-treated neutrophils. **(C)** Human neutrophil-like dHL-60 cells were either left untreated (Unt.) or treated with recombinant human LOX (75 ng/mL) ± NE inhibitor sivelestat (50 µM) for 6h before brief stimulation with fMLP (10 nM) and supernatant collection. Fluorometric analysis of the supernatant/EnzChek^™^ elastase substrate mix revealed that the LOX-induced ∼1.6-fold increase (p<0.001) in extracellular NE activity is blocked by sivelestat. **(D)** Flow cytometry-based analysis of FITC-Annexin V-labeled HRECs following 16h co-culture with LOX-pretreated or untreated (Unt.) dHL-60 cells in the absence or presence of NE inhibitor sivelestat (50 µM) revealed that LOX-treated dHL-60 cells cause a significant increase in HREC apoptosis, which is blocked in the presence of sivelestat. (**E**) Untreated (Unt.) or LOX-treated (75 ng/mL; 6h) dHL-60 cells were labelled with anti-CD63 (to visualize NE-containing granules; *green*) and Phalloidin-594 (to visualize F-actin; *red*). Representative confocal images and subsequent quantitative analysis, performed as depicted in the schematic, demonstrate that LOX causes a significant increase in co-localized CD63 aggregation and F-actin polarization (indicated by arrowheads in fluorescence images). Column scatter plots indicate mean and distribution from ≥80 cells. **(F)** AFM stiffness measurement of untreated (Unt.) or LOX-treated (75 ng/mL; 6h) dHL-60 cells (n=30) revealed that LOX induces significant softening (by ∼70%, p<0.001) of dHL-60 cells. Bar graphs indicate mean ± SEM. Scale bar, 5 µm.

We also looked at the effects of LOX on human differentiated HL-60 (dHL-60) cells that are commonly used as surrogates for primary human neutrophils (24–26). LOX exerted similar dose- and time-dependent effects on dHL-60 cells as on the mouse neutrophils, with the same 75 ng/mL dose producing a maximal increase in NE mRNA levels (by 1.25 fold; p<0.001) and extracellular NE activity (by ∼2 fold; p<0.05) in dHL-60 cells at 6h (**Supplementary** Fig. 4A, B). Notably, the LOX-induced increase in extracellular NE activity correlated with an ∼1.3-fold (p<0.001) increase in CD63 surface density (**Supplementary** Fig. 4C), presumably resulting from the fusion of CD63-rich azurophilic granules with cell membrane during NE release. Consistent with these findings, and similar to mouse neutrophils, dHL-60 cells treated with LOX exhibited a significant increase in extracellular NE activity and retinal EC apoptosis in co-culture studies (**Fig. 2C, D**). This retinal EC death was mediated by NE because addition of sivelestat, which inhibits extracellular NE activity (**Fig. 2C**), concomitantly blocked the dHL-60 cell-induced cytotoxicity towards HRECs (**Fig. 2D**).

Membrane accumulation of azurophilic granule marker CD63 is indicative of extracellular NE secretion from activated neutrophils (31, 32). Indeed, we observed that the higher extracellular NE activity in LOX-treated dHL-60 cells is associated with a significant increase (by 1.25 fold; p<0.05) in CD63 aggregation (**Fig. 2E**). Activated neutrophils are also known to generate higher levels of cytotoxic superoxide resulting from membrane assembly of cytosolic p47^phox^, a key organizer for NADPH oxidase machinery. Consistent with this, LOX-treated dHL-60 cells also exhibited a significant increase (by 1.3 fold; p<0.05) in p47^phox^ accumulation (**Supplementary** Fig. 5).

Past studies have reported F-actin remodeling in activated neutrophils (18, 19, 33). To determine whether LOX induces F-actin reorganization during neutrophil activation, we labeled LOX-treated dHL-60 cells with rhodamine-phalloidin. Fluorescence images and quantitative analysis revealed that LOX treatment significantly increases (by 1.3-fold; p<0.05) F-actin polarization, with higher F-actin density seen in regions of CD63 and p47^phox^ aggregation (**Fig 2E and Supplementary** Fig. 5). Interestingly, this LOX-induced F-actin polarization was associated with a significant decrease in dHL-60 cell stiffness (by 66%; p<0.001), as judged by our AFM measurements (**Fig. 2F**).

### LOX induces rapid and transient actin remodelling in neutrophils

F-actin remodeling is a recognized hallmark of activated neutrophils (18, 19, 33). To assess its precise role (stimulatory vs inhibitory) and spatiotemporal regulation during LOX-induced neutrophil activation, we systematically investigated the actin cytoskeletal dynamics in LOX-treated dHL-60 cells. As shown in **Fig. 3A**, LOX treatment resulted in rapid F-actin depolymerization (within 10 min), which led to a 65% decrease (p<0.001) in F-actin intensity. Interestingly, this rapid and substantial loss in F-actin was immediately followed by actin polymerization that increased up to 6 h when F-actin intensity increased by 1.8 fold (p<0.001) over the untreated condition.

**Figure 3:**
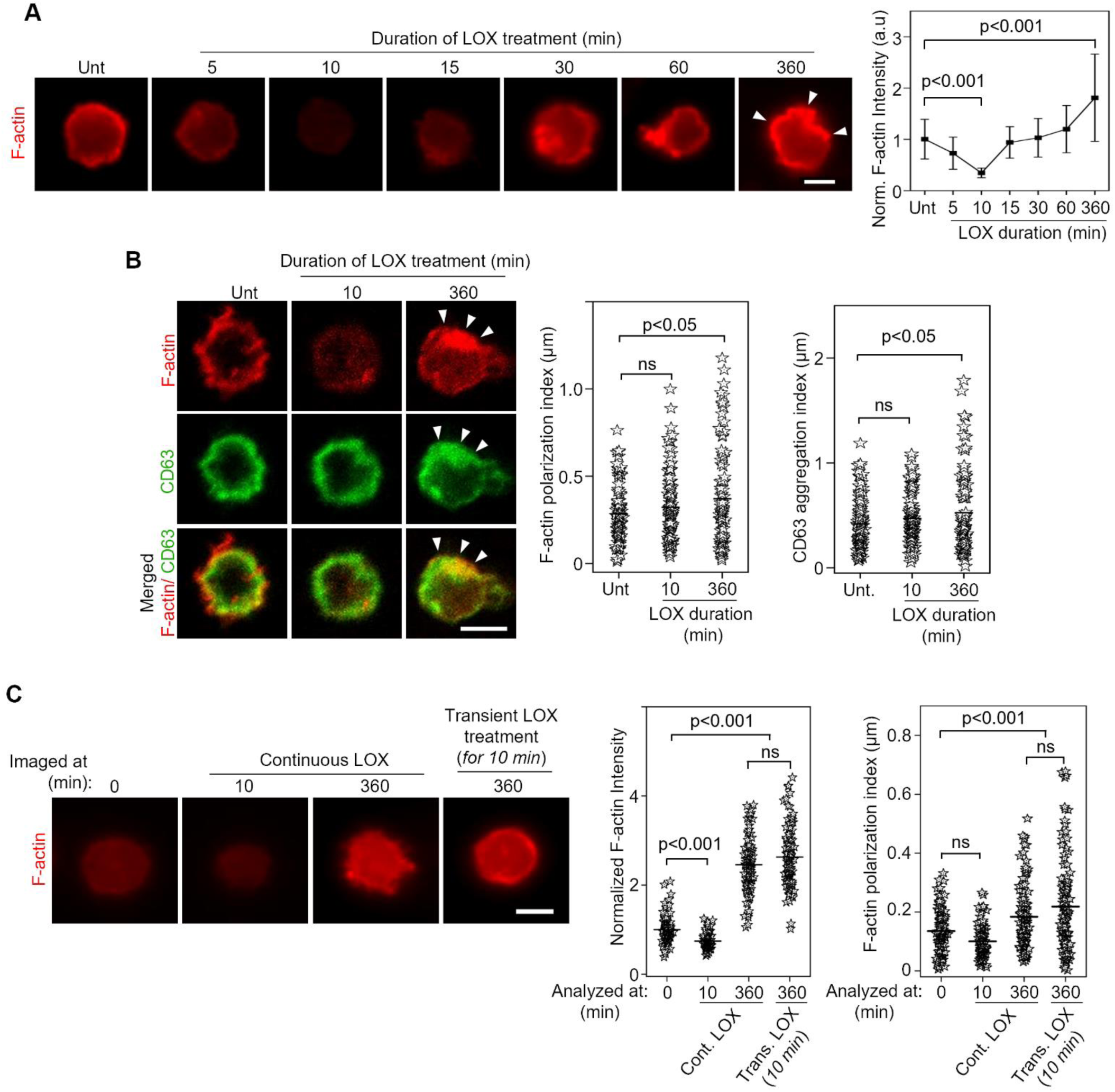
LOX induces rapid and transient actin remodelling in neutrophils. **(A)** dHL-60 cells were either left untreated (Unt.) or treated with LOX (75 ng/mL) for the indicated durations prior to labelling with Phalloidin-594 to visualize F-actin. Representative fluorescence images and subsequent quantitative analysis revealed that LOX causes a rapid and transient decrease in F-actin intensity followed by a significant increase at 360 min (6h). Line graph indicates mean ± SD from ≥200 cells; intensity was normalized with respect to untreated (Unt.) cells. **(B)** Untreated (Unt.) or LOX-treated (6h) dHL-60 cells were labelled with Phalloidin-594 (*red*) and anti-CD63 (to visualize NE-containing granules; *green*). Representative confocal images and subsequent quantitative analysis demonstrate that LOX-induced CD63 aggregation is spatiotemporally associated with F-actin polarization (arrowheads) at 360 min (6h). Plots indicate mean and distribution from ≥90 cells. **(C)** dHL-60 cells were treated with 75 ng/mL LOX either continuously for 360 min (6h) or transiently for 10 min prior to labelling with Phalloidin-594 at the indicated time points. Representative fluorescence images and subsequent quantitative analysis revealed that removal of LOX after 10 min does not inhibit F-actin repolymerization seen at 6h. Plots indicate mean and distribution from ≥110 cells; F-actin intensity was normalized with respect to value at 0 min. Bar graphs indicate mean ± SEM. Scale bar, 5 µm.

We also observed that while CD63 and p47^Phox^ remained uniformly distributed during rapid F-actin depolymerization (at 10 min), they underwent significant aggregation during the phase of actin re-polymerization and polarization (at 6h), co-localizing appreciably with the polarized F-actin (**Fig. 3B and Supplementary** Fig. 6A). Notably, this time-dependent effects of LOX on CD63 and p47^Phox^ distribution was mirrored in the extracellular release of NE and superoxide whose levels remained unchanged at 10 min but increased significantly at 6h (**Supplementary** Fig. 6B,C).

LOX induced a biphasic change in F-actin in dHL-60 cells, with the rapid depolymerization followed by a significant increase in actin polymerization and polarization. To determine whether LOX actively contributes to both phases of actin remodeling, dHL-60 cells were treated with LOX either transiently (for only 10 min) or continuously for the 6h duration prior to assessment of F-actin density and organization. As shown in **Fig. 3C**, the transient LOX treatment, where LOX was removed from dHL-60 culture medium after 10 min, led to a comparable increase in F-actin density (by ∼2.5 fold; p<0.001) and polarization (by ∼1.4 fold; p<0.001) at 6h as that seen with the continuous LOX treatment. Thus, LOX appears to contribute only to F-actin depolymerization, with the second phase of actin re-polymerization and polarization seemingly occurring in a LOX-independent manner.

### Transient actin depolymerization is necessary for LOX-induced neutrophil activation

LOX-induced increase in extracellular NE and superoxide release was associated with extensive actin remodeling in dHL-60 cells. To determine whether actin remodeling plays a causal role in LOX-induced dHL-60 cell activation, actin-stabilizing drug Jaspakinolide (JASP) (27) was added to dHL-60 cultures simultaneously with LOX. As shown in **Figs. 4A and 4B**, JASP blocked the LOX-induced rapid F-actin depolymerization, which also inhibited the subsequent actin re-polymerization and polarization seen over the 6h period. Thus, the early phase of (LOX-induced) actin depolymerization appears to be a prerequisite for late-phase (LOX-independent) remodeling.

**Figure 4:**
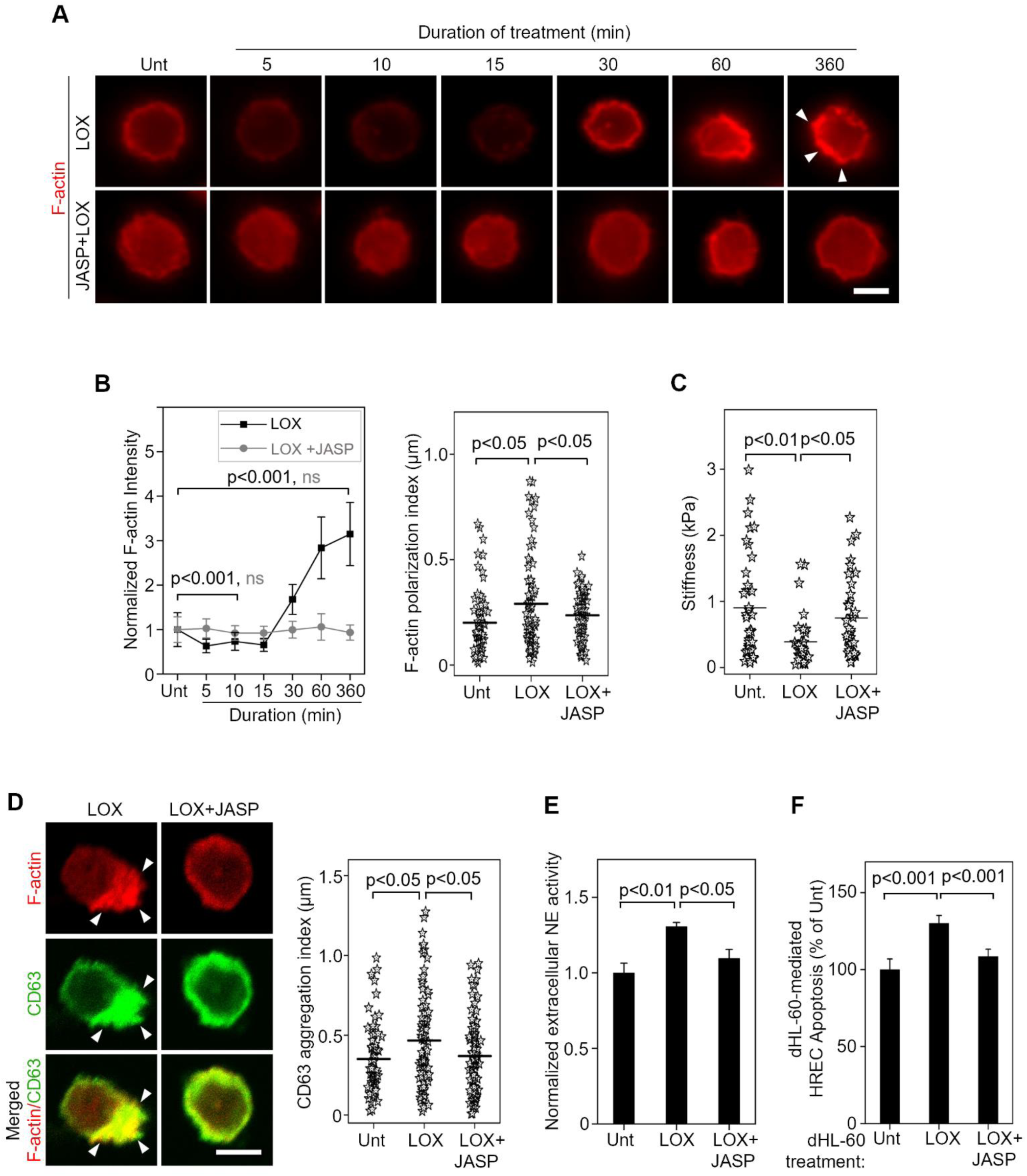
Transient actin depolymerization is necessary for LOX-induced neutrophil activation. **(A,B)** dHL-60 cells were treated with 75 ng/mL LOX ± actin stabilizing drug JASP (1 µM) for the indicated durations prior to labelling with Phalloidin-594 to visualize F-actin. Representative fluorescence images (A) and subsequent quantitative analysis (B) revealed that the LOX-induced F-actin remodeling is blocked by JASP. Line graph in (B) indicates mean ± SD from ≥80 cells; intensity was normalized with respect to untreated (Unt.) dHL-60 cells. Plots indicate mean and distribution from ≥65 cells. $p<0.001 for Unt Vs LOX treated cells; #p=ns for unt vs LOX+JASP treated cells. **(C)** AFM stiffness measurement of dHL-60 cells (n=40) that were either left untreated (Unt.) or treated with 75 ng/mL LOX ± actin stabilizing drug JASP (1 µM) for 6h revealed that the LOX-induced softening of dHL-60 cells is significantly inhibited (by 40%; p<0.05) by JASP. **(D)** Untreated (Unt.) or LOX ± JASP-treated (6h) dHL-60 cells were labelled with Phalloidin-594 (*red*) and anti-CD63 (to visualize NE-containing granules; *green*). Representative confocal images and subsequent quantitative analysis revealed that JASP treatment concurrently blocks LOX-induced F-actin polarization and CD63 aggregation (arrowheads). Plots indicate mean and distribution from ≥70 cells. **(E)** Untreated (Unt.) or LOX ± JASP-treated (6h) dHL-60 cells were briefly stimulated with fMLP (10 nM) prior to supernatant collection. Fluorometric analysis of the supernatant/EnzChek^™^ elastase substrate mix revealed that the LOX-induced increase in extracellular NE activity is significantly inhibited (by ∼70%; p<0.05) by JASP. Unt: Untreated. **(F)** HRECs were co-cultured with [LOX±JASP]-pretreated or untreated (Unt.) dHL-60 cells for 16h prior to FITC Annexin V/PI labeling and flow cytometry. Quantification of the total number of Annexin V-positive HRECs (plotted as % of Unt.) revealed that the increase in HREC apoptosis caused by LOX-pretreated dHL-60 cells is significantly inhibited (by 70%; p<0.001) by JASP. Bar graphs indicate mean ± SEM. Scale bar, 5 µm.

In addition to preventing F-actin remodeling, JASP also inhibited the LOX-induced softening of dHL-60 cells (**Fig. 4C**), which indicates a causal link between LOX and actin-dependent neutrophil stiffness. Further, these inhibitory effects of JASP on LOX-induced changes in dHL-60 cell mechanics were associated with significant reduction in the aggregation of CD63 (**Fig. 4D**) and p47^Phox^ (**Supplementary** Fig. 7A) that, in turn, led to a >70% inhibition (p<0.05) in LOX-induced increase in extracellular NE activity (**Fig. 4E**) and superoxide generation (**Supplementary** Fig. 7B), respectively. Predictably, this suppression of LOX-induced dHL-60 cell activation resulted in a 70% inhibition (p<0.001) in dHL-60 cell cytotoxicity towards HRECs (**Fig. 4F**). This finding indicates that LOX-mediated actin depolymerization is required for dHL-60 cell activation and cytotoxicity.

### Transient actin depolymerization is sufficient to cause neutrophil activation and cytotoxicity

Since we found that early F-actin depolymerization is necessary for LOX-induced dHL-60 cell activation and cytotoxicity, we next asked whether F-actin depolymerization alone is sufficient to activate dHL-60 cells. To address this, cells were treated with the F-actin disrupting agent cytochalasin D (cytoD; 2.5 µM) for 6h. As expected, cytoD treatment led to rapid F-actin depolymerization in dHL-60 cells, resulting in ∼80% inhibition (p<0.001) in F-actin density (**Fig. 5A**). Notably, this rapid loss of F-actin was followed by a significant increase in actin polymerization (by ∼1.15 fold; p<0.001) and polarization (by ∼1.35 fold; p<0.05) over the untreated condition (**Fig. 5B**), thus mirroring the biphasic actin remodeling seen in response to LOX treatment.

**Figure 5:**
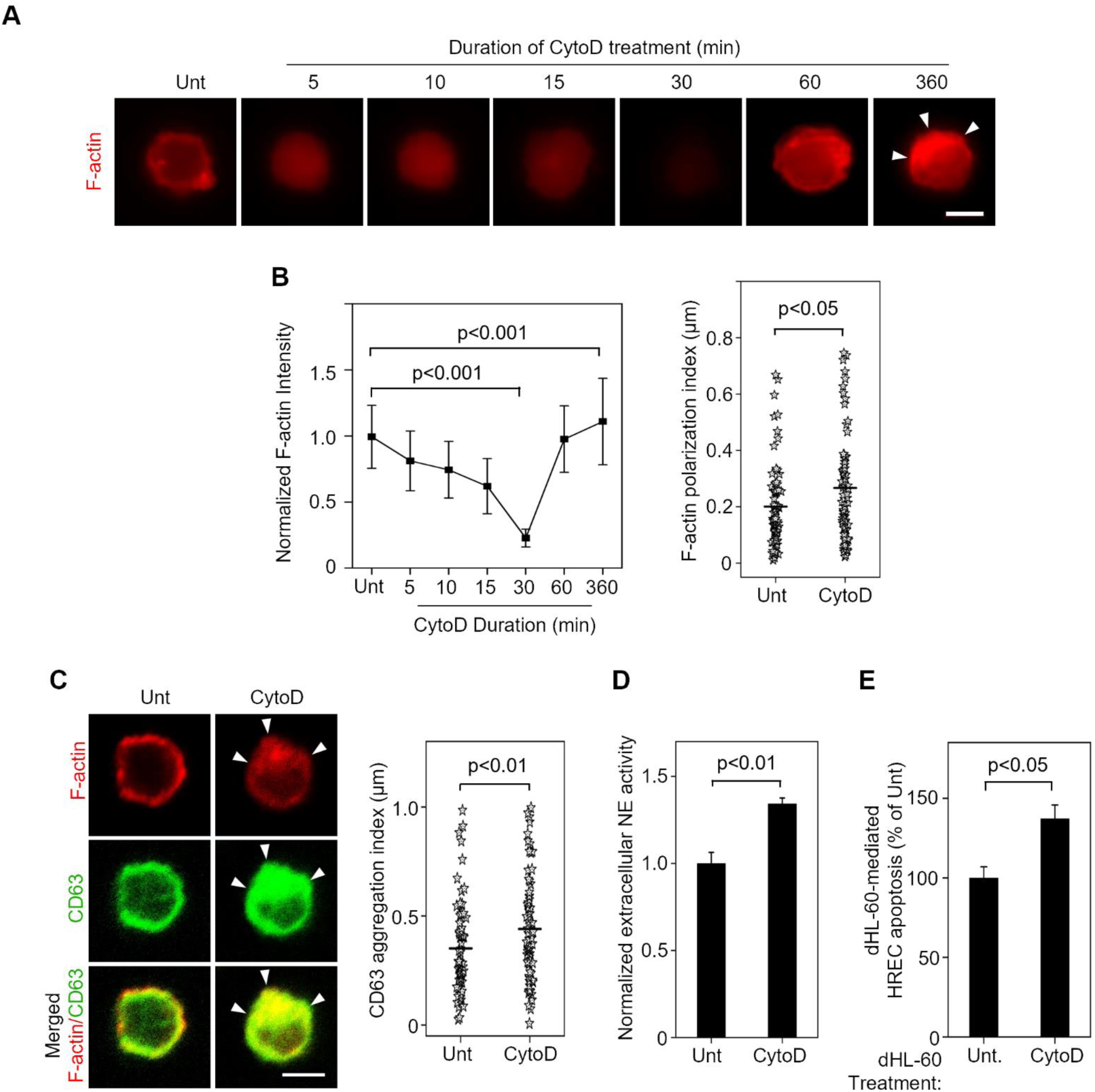
Transient actin depolymerization is sufficient to cause neutrophil activation and cytotoxicity. **(A, B)** dHL-60 cells were either left untreated (Unt.) or treated with actin depolymerization drug cytoD (2.5 µM) for the indicated durations prior to labelling with Phalloidin-594 to visualize F-actin. Representative fluorescence images and subsequent quantitative analysis revealed that cytoD causes a transient decrease in F-actin intensity followed by a significant increase in F-actin polymerization and polarization at 360 min (6h). Line graph indicates mean ± SD from ≥200 cells; intensity was normalized with respect to untreated (Unt.) cells. Plots indicate mean and distribution from ≥70 cells. **(C)** Untreated (Unt.) or cytoD-treated (6h) dHL-60 cells were labelled with Phalloidin-594 (*red*) and anti-CD63 (to visualize NE-containing granules; *green*). Representative confocal images and subsequent quantitative analysis revealed that cytoD treatment leads to a significant increase in co-localized F-actin polarization and CD63 aggregation (arrowheads) at 360 min (6h). Plots indicate mean and distribution from ≥65 cells. **(D)** Untreated (Unt.) or cytoD-treated (6h) dHL-60 cells were briefly stimulated with fMLP (10 nM) prior to supernatant collection. Fluorometric analysis of the supernatant/EnzChek^™^ elastase substrate mix revealed that cytoD-treated dHL-60 cells exhibit a significant increase (by 30%; p<0.01) in extracellular NE activity when compared with untreated (Unt.) cells. **(E)** HRECs were co-cultured with cytoD-pretreated or untreated (Unt.) dHL-60 cells for 16h prior to FITC Annexin V/PI labeling and flow cytometry. Quantification of the total number of Annexin V-positive HRECs (plotted as % of Unt.) revealed that cytoD-treated dHL-60 cells cause a significant increase (by ∼40%; p<0.05) in HREC apoptosis. Bar graphs indicate mean ± SEM. Scale bar, 5 µm.

Further, and similar to LOX-treated cells, those treated with cytoD exhibited a significant increase in the aggregation of CD63 (by 1.25 fold; p<0.01) and p47^Phox^ (by 1.3-fold; p<0.01) during the 6h period of actin repolymerization and polarization, co-localizing appreciably with the polarized F-actin (**Fig. 5C and Supplementary** Fig. 8A). Predictably, this cytoD-induced CD63 and p47^Phox^ aggregation in dHL-60 cells led to a concurrent increase in extracellular NE activity (by 1.25 fold; p<0.01) (**Fig. 5D**), superoxide generation (by 1.45 fold; p<0.01) (**Supplementary** Fig. 8B), and cytotoxicity towards HRECs (by 1.45 fold; p<0.05) (**Fig. 5E**).

### LOX inhibition blocks diabetes-induced F-actin remodeling and CD63 aggregation in neutrophils

Our aforementioned findings indicate that F-actin remodelling is essential for LOX-induced activation and cytotoxicity of dHL-60 cells. Since we previously showed (Fig. 1) that LOX inhibition rescues mouse neutrophil activation and cytotoxicity in diabetes, we asked whether this protective effect of LOX inhibition results from prevention of F-actin remodelling and associated CD63/p47^Phox^ aggregation. Indeed, phalloidin labelling of mouse neutrophils revealed that diabetes leads to a significant increase in F-actin density (by ∼1.3 fold; p<0.001) and polarization (by 1.25 fold; p<0.001) that are blocked by LOX inhibitor BAPN (**Fig. 6A,B**). The same trend was observed for the aggregation of CD63 (**Fig. 6A,B**) and p47^Phox^ (**Supplementary** Fig. 9) that increased significantly in diabetes but were suppressed by LOX inhibition. Finally, our AFM measurements revealed that the diabetes-induced F-actin remodeling in mouse neutrophils is associated with a 40% decrease (p<0.01) in cell stiffness (**Fig. 6C**), which is also significantly rescued by LOX inhibition.

**Figure 6:**
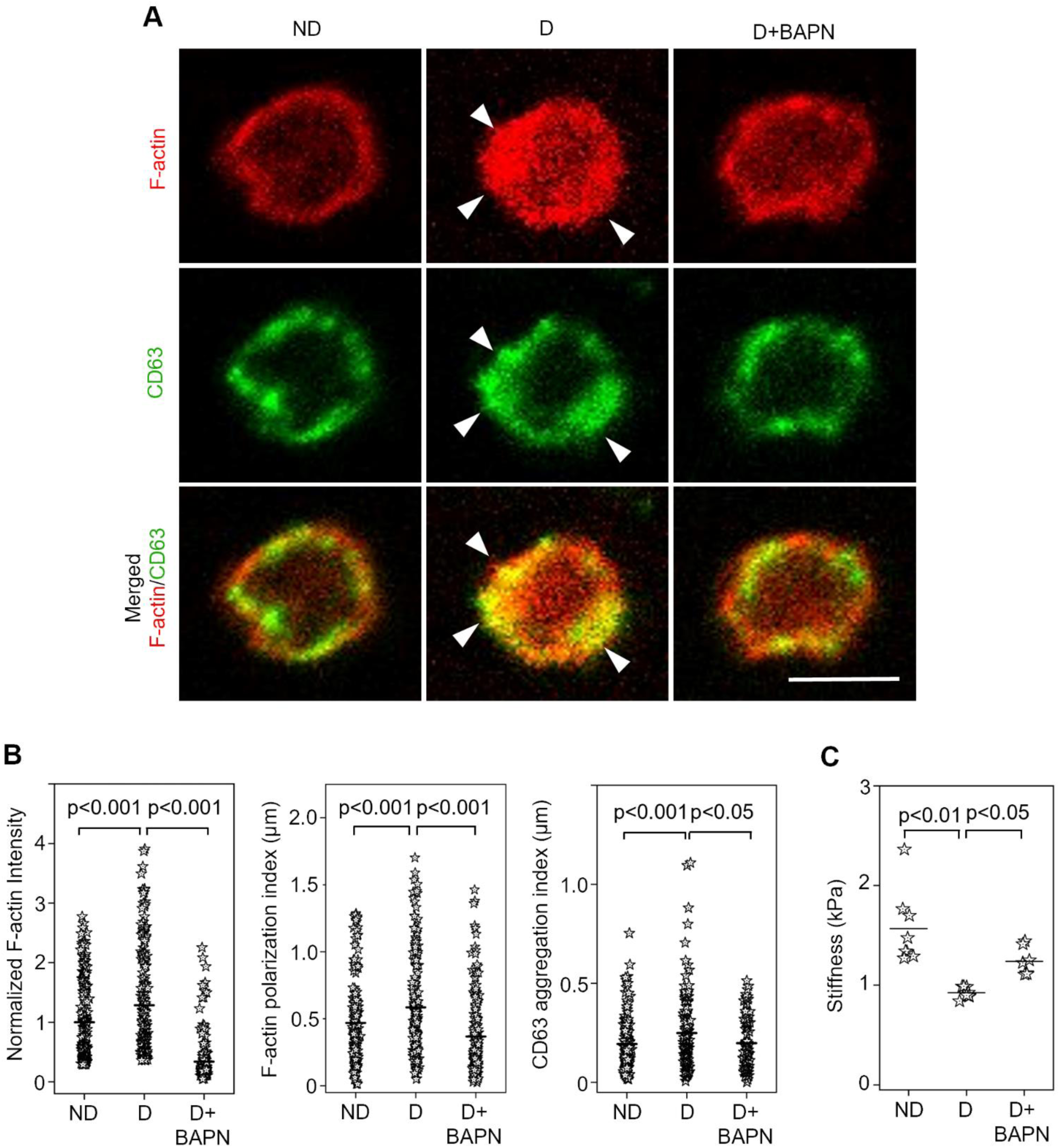
LOX inhibition blocks diabetes-induced F-actin remodeling and CD63 aggregation in neutrophils. Mouse bone marrow neutrophils isolated from nondiabetic (ND), diabetic (D), or D mice treated with BAPN (D+BAPN; 3 mg/kg BW) (n≥90 cells/group; 10 wk diabetes) were labelled with Phalloidin-594 (to visualize F-actin; *red*) and anti-CD63 (to visualize NE-containing granules; *green*). **(A)** Representative confocal images and **(B)** subsequent quantitative analysis revealed that LOX inhibition using BAPN prevented the diabetes-induced concurrent increase in F-actin intensity, F-actin polarization, and CD63 aggregation. Arrowheads indicate F-actin polarization or CD63 aggregation. Plots indicate mean and distribution from ≥90 cells. Scale bar, 5 µm. **(C)** AFM stiffness measurement of bone marrow neutrophils (n≥15 cells) isolated from nondiabetic (ND), diabetic (D), or D mice treated with BAPN (D+BAPN; 3 mg/kg BW) (n=6 mice/group; 10 wk diabetes) revealed that the diabetes-induced reduction in neutrophil stiffness (by 41%; p<0.01) is significantly inhibited (by 48%; p<0.05) by LOX inhibitor BAPN.

## DISCUSSION

Past studies have implicated neutrophil activation (release of NE and ROS) in retinal EC apoptosis and capillary degeneration associated with early DR (3, 4). But precisely how neutrophils become activated in diabetes is not fully understood. The current study addresses this knowledge gap by showing that LOX, which is upregulated in the retina in diabetes, directly activates neutrophils and increases their cytotoxicity towards retinal ECs. Specifically, we showed that LOX induces F-actin remodelling and cell softening in neutrophils that, in turn, facilitates the aggregation of CD63-containing granules and NADPH organizer p47^Phox^, thereby leading to increased extracellular release of cytotoxic NE and superoxide that cause retinal EC apoptosis (**Figure 7**). This finding is important as it highlights a *dual* proinflammatory role for LOX in early DR wherein it activates neutrophils in its soluble form while simultaneously increasing EC susceptibility to neutrophil-induced cytotoxicity in its matrix-localized form (7). Thus, LOX promises to be a potent anti-inflammatory target for clinical DR treatment.

**Figure 7:**
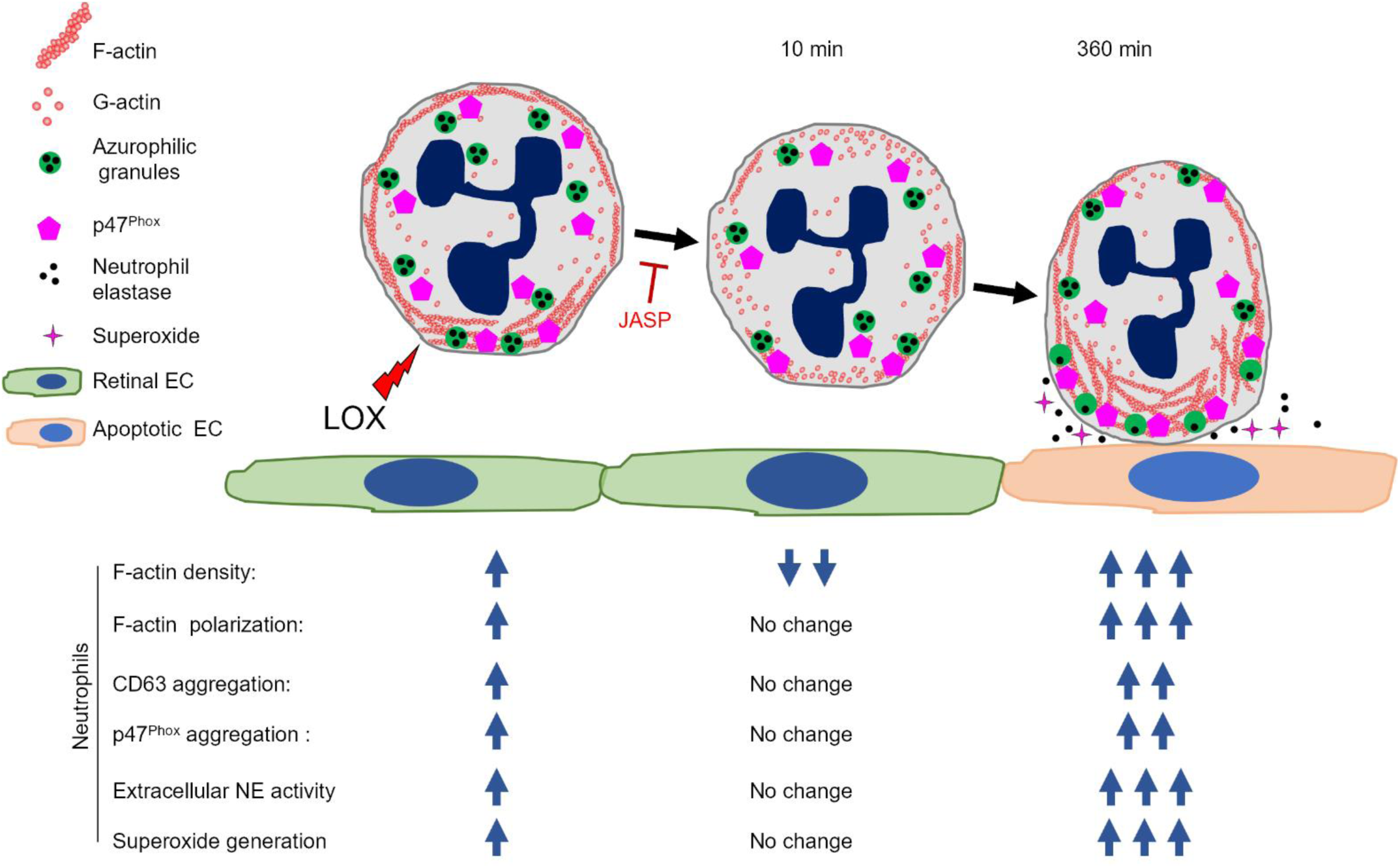
Schematic illustration of LOX-induced regulation of neutrophil actin dynamics, activation, and cytotoxicity in diabetes. Based on our current findings, we propose that LOX is a mechanical determinant of neutrophil activation wherein it alters actin cytoskeleton and cell stiffness to increase extracellular NE and superoxide release that, together, exert cytotoxic effects on retinal ECs. This newly identified mechanism of neutrophil activation begins with LOX-induced rapid (within ∼10 min) F-actin depolymerization. At this early time, the membrane distribution of CD63 and p47^Phox^, which reflects the cell’s ability to release extracellular NE and superoxide, respectively, remains uniform and unchanged. However, over time (6h), as F-actin recovers through repolymerization, it simultaneously becomes polarized. Interestingly, this late-phase F-actin remodeling causes redistribution (aggregation) of CD63 and p47^Phox^ that predictably leads to increased extracellular release of NE and superoxide, key cytotoxic mediators of retinal EC death in diabetes.

The biological effects of LOX have been commonly associated with its matrix-localized form that crosslinks collagen/elastin and stiffens extracellular matrix (9, 10). Supporting this view, our recent studies revealed that increased levels of retinal subendothelial matrix LOX increases capillary stiffness in diabetes that, in turn, contributes to retinal vascular inflammation and capillary degeneration associated with early DR (7). This mechanical regulation of retinal capillary loss is attributed to the activation of mechanosensitive NF-κB in retinal ECs, which leads to endothelial RAGE and ICAM-1 upregulation, increased neutrophil adhesion, and greater EC susceptibility to neutrophil/NE-induced cytotoxicity (6–8). However, LOX and its family members LOX-like proteins 1-4 (LOXL1-4) also exist in an alternate soluble form that can induce intracellular changes at the transcriptional, epigenetic, and cytoskeletal level (9, 14). Specifically in cancer cells, such LOX-induced intracellular changes have been implicated in EMT that drives tumor progression and metastasis (9, 34). Whether and how soluble LOX alters immune cell function is, however, poorly understood. Given that LOX inhibition in diabetic mice reduces leukocyte-EC adhesion and retinal EC death, we asked whether soluble LOX could directly activate neutrophils and make them cytotoxic towards retinal ECs.

Indeed, we found that treatment of mouse neutrophils or dHL-60 cells with recombinant LOX significantly enhances their activation (increased NE and superoxide release) and cytotoxicity towards retinal ECs. Further, and perhaps more importantly, the characteristic neutrophil activation and cytotoxicity towards retinal ECs observed in 10 week-diabetic mice was blocked by pharmacological LOX inhibition. Collectively, these findings indicate that LOX plays a causal role in diabetes-induced neutrophil activation and cytotoxicity. To our knowledge, such a direct effect of LOX on neutrophils, or leukocytes in general, has not been reported before. Additionally, we noted that (a) neutrophils remain quiescent in short-term (four weeks’ duration) diabetic mice that do not exhibit retinal LOX upregulation, and (b) LOX levels do not increase in neutrophils isolated from 10-wk diabetic mice. These findings support the idea that neutrophil activation in diabetes is caused by an increase in exogenous (retina-derived), rather than endogenous (neutrophil-derived), LOX. Interestingly, although we did not observe neutrophil LOX upregulation in diabetes, a recent study demonstrated an increase in LOXL-4 expression in circulating and tumor-infiltrating neutrophils of individuals with colorectal cancer liver metastases that are resistant to anti-angiogenic therapy (35). Whether the increase in neutrophil LOXL-4 regulated neutrophil function within the tumor microenvironment was, however, not assessed.

NE released by activated neutrophils contributes significantly to retinal EC apoptosis in diabetes (3). Consistent with this, we found that addition of NE inhibitor sivelestat blocked the LOX-induced neutrophil cytotoxicity towards retinal ECs, thus confirming that LOX-induced increase in NE contributed to retinal EC death. NE is stored within cytosolic azurophilic granules that contain an integral membrane protein CD63. In stimulated neutrophils, these granules aggregate and fuse with the plasma membrane, leading to an increase in CD63 membrane density and extracellular NE release (31, 36, 37). CD63 has also been implicated in the processing of pro-NE to mature NE (38). Predictably, our findings revealed that the LOX-induced increase in NE release was associated with significant aggregation and membrane deposition of CD63 immunolabeled granules. Notably, the LOX-induced NE release and CD63 aggregation were spatiotemporally associated with increased superoxide production and clustering of p47^Phox^, a cytosolic protein that assembles at membrane sites of activated neutrophils to functionally organize the superoxide-generating NADPH oxidase (39, 40). This ability of LOX to facilitate the release of both NE and superoxide, which involve distinct molecular players and signaling pathways, underscores its importance in regulating neutrophil activation.

To understand how LOX might promote CD63 and p47^phox^ aggregation associated with NE and superoxide release, respectively, we looked at the role of actin cytoskeletal remodeling that is implicated in granule trafficking and NADPH oxidase assembly in neutrophils (15–17). The important role of actin cytoskeleton in regulating neutrophil physiology can be further appreciated from studies that have linked defects in its remodeling to severe neutrophil dysfunction, as seen in “lazy-leukocyte syndrome” (41, 42). Actin remodeling is a dynamic process that involves the assembly (polymerization) and disassembly (depolymerization) of F-actin, which forms a cortical filamentous layer adjacent to the neutrophil plasma membrane. The cortical actin is believed to form a barrier that prevents granule access to plasma membrane because treatments or proinflammatory factors (e.g. TNFα) that depolymerize actin lead to increased neutrophil exocytosis (15, 16, 18, 43, 44). Thus, we looked to see if the LOX-induced CD63 and p47^phox^ aggregation and NE/superoxide release in neutrophils/dHL-60 cells are mediated by actin remodeling.

Our findings revealed that LOX treatment causes rapid F-actin depolymerization (within 10 min) in dHL-60 cells, followed by gradual but extensive re-polymerization. Such rapid actin depolymerization has also been seen in TNFα-treated neutrophils (18). Notably, removing LOX after the rapid depolymerization phase did not impair actin re-polymerization. Thus, LOX does not appear to contribute to the second phase of actin recovery. We also observed that actin re-polymerization resulted in a significant increase in F-actin density (above the initial levels), reflecting overcompensation during recovery. Interestingly, re-polymerization of actin (over 6h) was accompanied by extensive polarization, which was spatiotemporally associated with CD63 and p47^Phox^ aggregation and a significant increase in extracellular NE and superoxide release. As these markers of dHL-60 activation remained unchanged during the early phase of rapid actin depolymerization, we believe that the gradually re-polymerizing actin (during recovery phase) orchestrates granule trafficking and NADPH oxidase organization, leading to their deposition/stabilization in the plasma membrane where they facilitate the release of NE and superoxide, respectively. This idea is consistent with previous findings in neutrophils that revealed (a) a direct crosstalk between the molecular regulators of actin remodeling and azurophilic granule secretory machinery (16), and (b) a direct interaction between actin and p47^Phox^ that stabilizes the NADPH oxidase machinery in plasma membrane (17).

To determine whether actin remodeling plays a causal role in LOX-induced CD63/p47^Phox^ aggregation and NE/superoxide release, we added actin-stabilizing agent jasplakinolide (JASP) to LOX-treated dHL-60 cells (27). Not only did JASP block the LOX-induced rapid actin depolymerization, it also prevented the subsequent actin re-polymerization and polarization as well as the concomitant aggregation of CD63 and p47^Phox^, NE and superoxide release, and cytotoxicity towards retinal ECs. Thus, actin remodeling appears to be an essential mediator of LOX-induced neutrophil activation, which is consistent with its role in TNFα-induced neutrophil activation (44). This ability of LOX to induce actin remodeling also extends to adherent cells, specifically cancer and smooth muscle cells (45, 46). That actin remodeling is sufficient to induce neutrophil activation was confirmed when cytoD treatment, which leads to rapid loss of F-actin, resulted in subsequent actin re-polymerization and polarization, CD63 and p47^Phox^ aggregation, NE and superoxide release, and cytotoxicity towards retinal ECs. Notably, these effects of JASP and cytoD on mouse neutrophil activation are mirrored in human neutrophils (43).

To assess the translational implications of these findings, we looked at actin remodeling in neutrophils isolated from 10 week-diabetic mice. As we showed in the current study, these mice exhibit retinal LOX upregulation that contributes to neutrophil activation (greater NE and superoxide release) and cytotoxicity towards retinal ECs. Predictably, we found that, similar to LOX-treated mouse neutrophils and dHL-60 cells, activated neutrophils from diabetic mice exhibit a significant increase in F-actin density and polarization along with CD63 and p47^Phox^ aggregation. Importantly, these diabetes-associated cytoskeletal, molecular, and functional changes in mouse neutrophils were blocked by pharmacological LOX inhibition. These findings in mouse neutrophils are in line with the observations of human neutrophils in type 2 diabetes where impaired actin polymerization is implicated in reduced exocytosis of primary granules, which is required for extracellular NE release (33). Thus, LOX-induced actin remodeling, characterized by rapid depolymerization followed by extensive re-polymerization, is believed to play a crucial role in neutrophil activation and cytotoxicity associated with diabetes.

Another notable effect of LOX on neutrophils was a significant decrease in cell stiffness. This LOX-induced cell softening can be attributed to actin remodeling because treatment with actin stabilizing agent JASP prevented the LOX-induced decrease in neutrophil stiffness. While we did not directly assess the implications of neutrophil stiffness in retinal EC dysfunction, we speculate that the increased flexibility resulting from cell softening facilitates the spreading of bound neutrophils on retinal ECs and helps them squeeze through the endothelial junctions (extravasation) for subsequent perivascular retinal damage. Future studies will be needed to elucidate the underlying intracellular mechanisms and pathological implications of LOX-induced mechanical changes in neutrophils.

By demonstrating the ability of soluble LOX to promote neutrophil activation and cytotoxicity in diabetes, the current findings offer a fresh perspective on the proinflammatory role of LOX in early DR that goes beyond its canonical matrix-stiffening effects. Our findings also imply that LOX exerts synergistic proinflammatory effects on the retinal vasculature through its soluble and matrix-localized forms that differentially activate neutrophils and increase retinal EC susceptibility to neutrophil-induced cytotoxicity, respectively. Thus, LOX appears to be a key proinflammatory stressor and a potential anti-inflammatory target for early DR. A deeper understanding of the mechanistic link between LOX, actin remodeling, and neutrophil activation may help identify new mechanobiology-based therapeutic targets for effective DR management in the future.

## Acknowledgement

This work was supported by National Institutes of Health, National Eye Institute grants R01EY028242 (to K.G.), R01EY033002, R01EY022938, and R24EY024864 (to T.S.K.), EY022938-S1 (to E.M.L.), and P30EY034070-01 (National Eye Institute core grant to Center for Translational Vision Research at the University of California, Irvine), and W.M. Keck Foundation Stephen Ryan Initiative for Macular Research (RIMR) Special Grant (to Doheny Eye Institute). This work was also supported by Research to Prevent Blindness, Inc., Unrestricted Grants to the Department of Ophthalmology at the University of California, Los Angeles and Gavin Herbert Eye Institute at the University of California, Irvine.

## Author Contributions

M.A. conceived the idea, designed and performed experiments, analyzed data, and wrote the manuscript, S.C. performed experiments and analyzed data. I.S.T. analyzed data, E.M.L. performed experiments and Z.R.B. performed experiments, T.S.K. designed experiments and analyzed data, K.G. conceived the idea, designed experiments, analyzed data, and wrote the manuscript. All authors reviewed, edited, and approved of the manuscript. K.G. is the guarantor of this work and, as such, has full access to all the data in the study and takes responsibility for the integrity of the data and the accuracy of the data analysis.

