## Supplementary Data for "Lysyl oxidase promotes actin-dependent neutrophil activation and cytotoxicity in diabetes: Implications for diabetic retinopathy"

**Supplementary Figure 1**

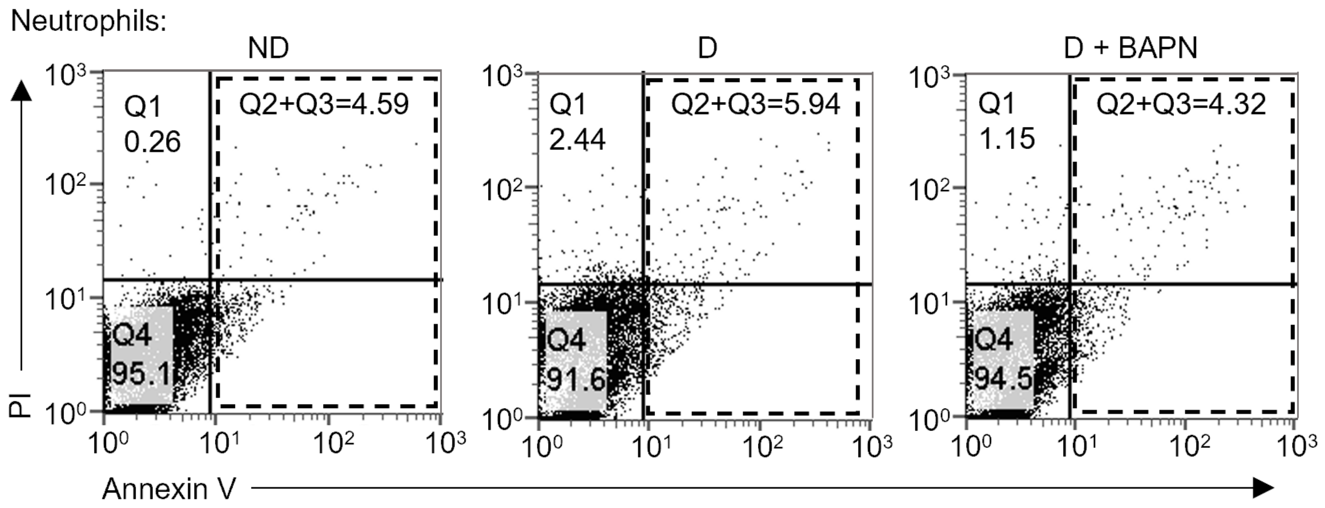


**Supplementary Figure 1: LOX promotes mouse neutrophil cytotoxicity towards retinal ECs in diabetes.** Representative flow cytometry scatter plots show FITC Annexin V- and PI-labeled mRECs following 16h of co-culture with mouse bone marrow-derived neutrophils isolated from nondiabetic (ND), diabetic (D), or D mice treated with LOX inhibitor BAPN (D+BAPN; 3 mg/kg BW) (n=6 mice/group; 10 weeks of diabetes). Annexin V-positive cells (Q2+Q3; dashed rectangle) indicates apoptotic mRECs following neutrophil co-culture.

**Supplementary Figure 2**

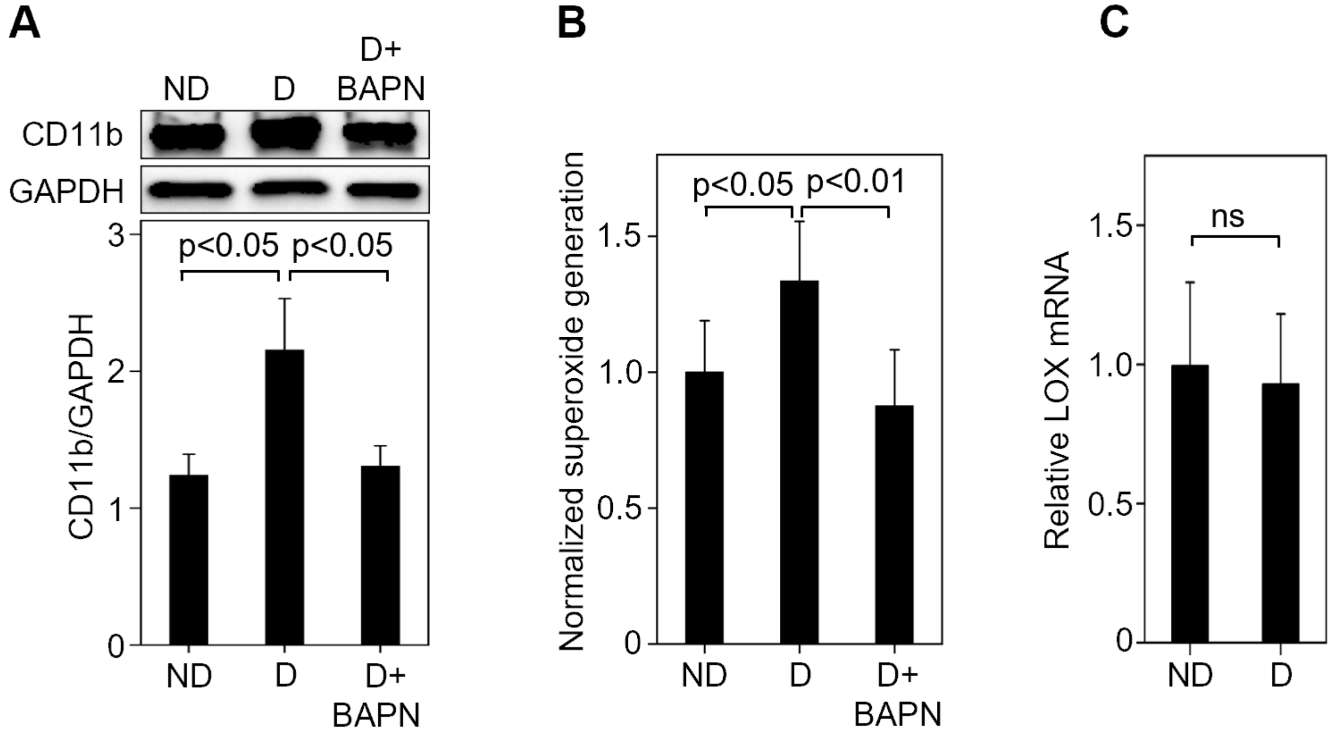


**Supplementary Fig 2: LOX contributes to diabetes-induced neutrophil activation. (A)** Representative Western blot bands and cumulative densitometric analysis of mouse neutrophils isolated from nondiabetic (ND), diabetic (D), or D mice treated with LOX inhibitor BAPN (D+BAPN; 3 mg/kg BW) (n=6 mice/group; 10 weeks of diabetes) indicate that the diabetes-induced 1.75-fold increase ($p<0.05$) in CD11b protein expression is prevented by LOX inhibition. **(B)** Neutrophils isolated from ND, D, or D+BAPN mice (n=6/group) were suspended in DHE solution dye before a brief stimulation with fMLP (10 nM). Fluorometric analysis revealed that the diabetes-induced ~35% increase ($p<0.05$) in superoxide generation is prevented by LOX inhibition. **(C)** RT-qPCR analysis of mouse neutrophils isolated from ND and D mice (n≥5 mice/group; 10 wk diabetes) indicate that diabetes does not alter LOX mRNA levels in neutrophils. Bar graphs indicate mean ± SD.

**Supplementary Figure 3**

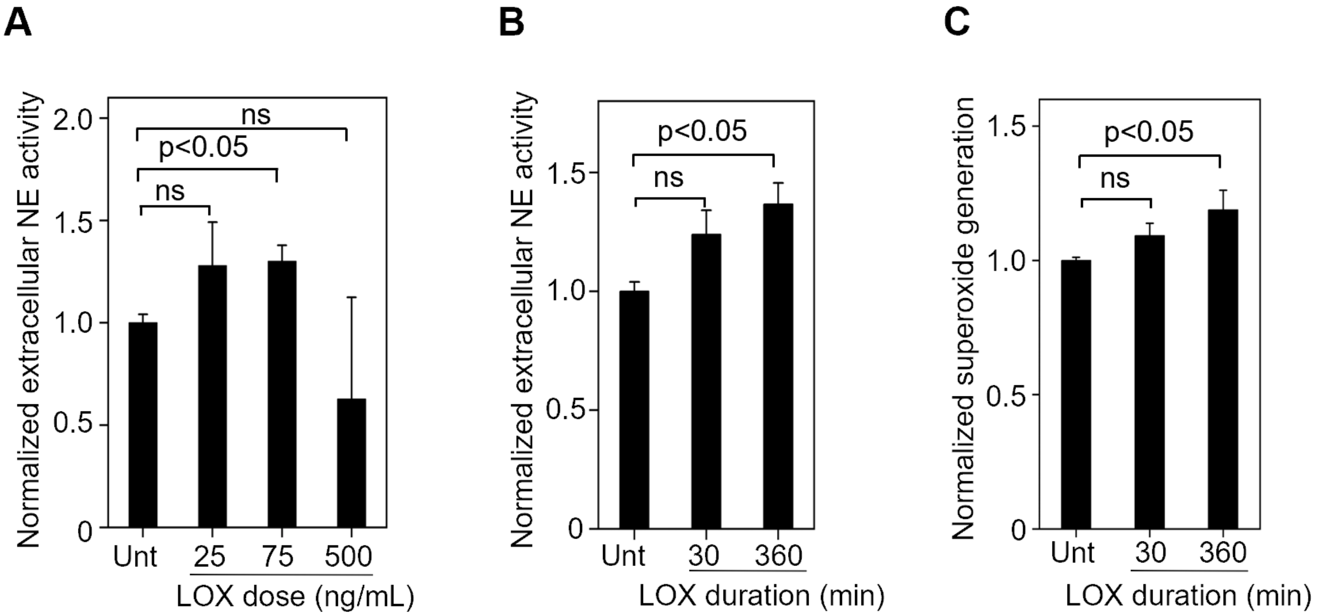


**Supplementary Fig 3: Dose- and time-dependent mouse neutrophil activation by LOX. (A)** Neutrophils isolated from ND mice and treated with recombinant mouse LOX for the indicated doses were briefly stimulated with fMLP (10 nM) prior to supernatant collection. Fluorescence intensity analysis of the supernatant/EnzChek™ elastase substrate mix revealed that 25-75 ng/mL LOX induces maximal extracellular NE activity. Bar graphs indicate mean ± SD. **(B)** Neutrophils isolated from ND mice and treated with 75 ng/mL recombinant LOX for the indicated durations were briefly stimulated with fMLP (10 nM) prior to supernatant collection. Fluorescence intensity analysis of the supernatant/EnzChek™ elastase substrate mix revealed that LOX induces a significant increase in extracellular NE activity at 360 min (6h). Bar graphs indicate mean ± SD. **(C)** Neutrophils isolated from ND mice were treated with 75 ng/mL recombinant LOX for the indicated durations prior to addition of superoxide indicator DHE and brief stimulation with fMLP (10 nM). Fluorometric analysis revealed that LOX induces a significant increase in superoxide generation at 360 min (6h). Bar graphs indicate mean ± SEM.

**Supplementary Figure 4**

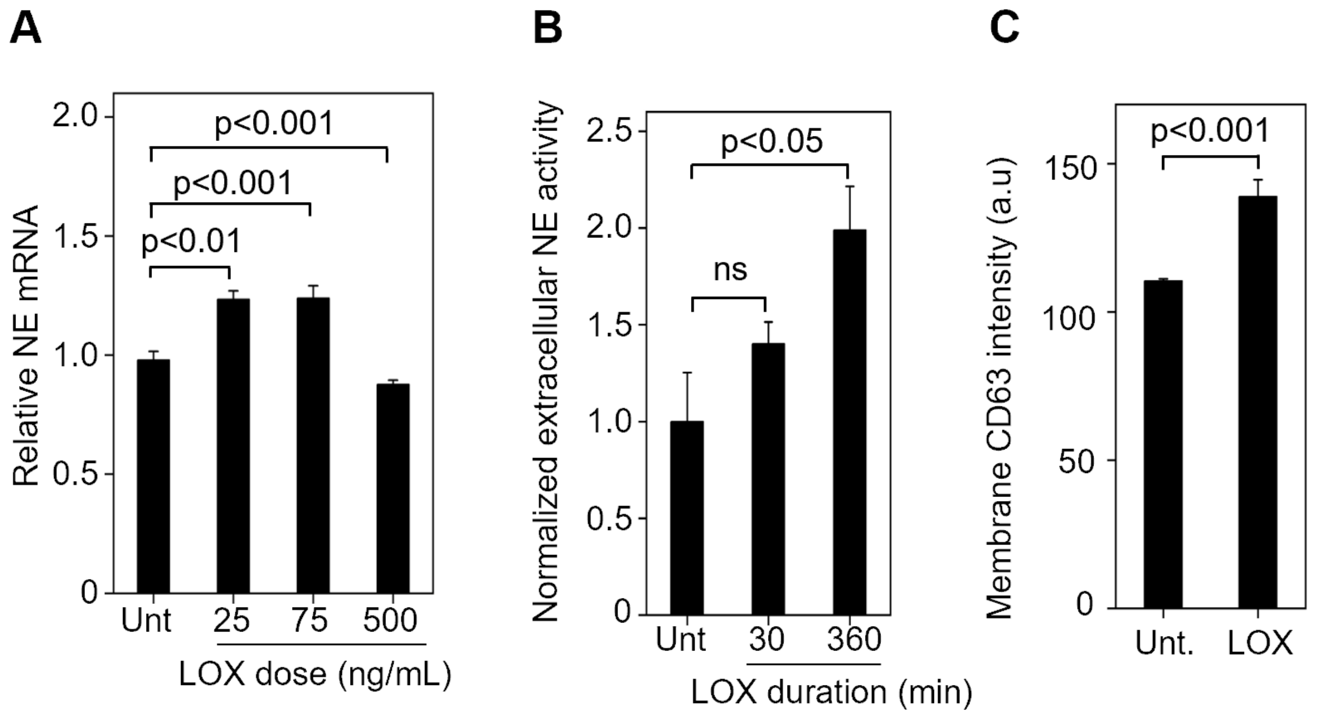


**Supplementary Fig 4: LOX activates dHL-60 cells. (A)** RT-qPCR analysis of dHL-60 cells treated with the indicated doses of recombinant human LOX for 360 min (6h) revealed that a dose of 25-75 ng/mL induces maximal NE mRNA expression. **(B)** dHL-60 cells treated with 75 ng/mL LOX for 30 or 360 min were briefly stimulated with fMLP (10 nM) prior to supernatant collection. Fluorescence intensity analysis of the supernatant/EnzChek™ elastase substrate mix revealed that LOX induces a significant increase in extracellular NE activity at 360 min (6h). Bar graphs indicate mean ± SD. **(C)** Flow cytometry-based analysis of dHL-60 cells treated with LOX (75 ng/ml; 6h) and labeled with anti-CD63 revealed that LOX induces a significant increase in CD63 surface density. Bar graphs indicate mean ± SEM.

**Supplementary Figure 5**

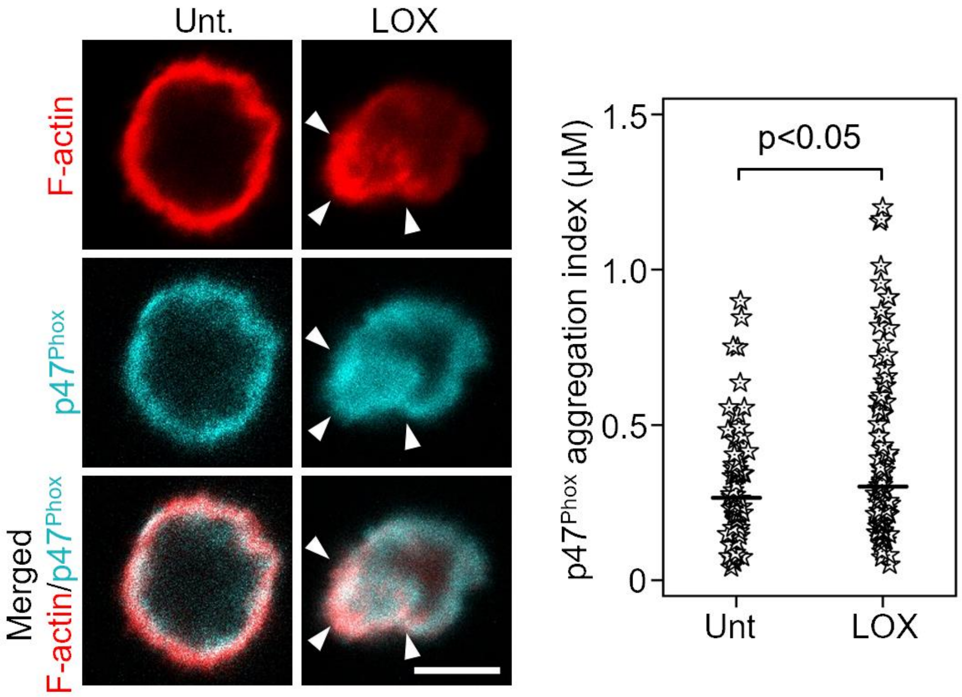


**Supplementary Fig 5: LOX induces aggregation of the NADPH organizer p47$^{phox}$.** Untreated (Unt.) or LOX-treated (75 ng/mL; 6h) dHL-60 cells were labelled with Phalloidin-594 (to visualize F-actin; *red*) and anti-p47$^{phox}$ (to visualize assembly of NADPH complex; *cyan*). Representative confocal images and subsequent quantitative analysis demonstrate that LOX induces co-localized F-actin polarization and p47$^{phox}$ aggregation (indicated by arrowheads). Column scatter plot indicates mean and distribution from ≥65 cells. Scale bar, 5 μm.

### Supplementary Figure 6

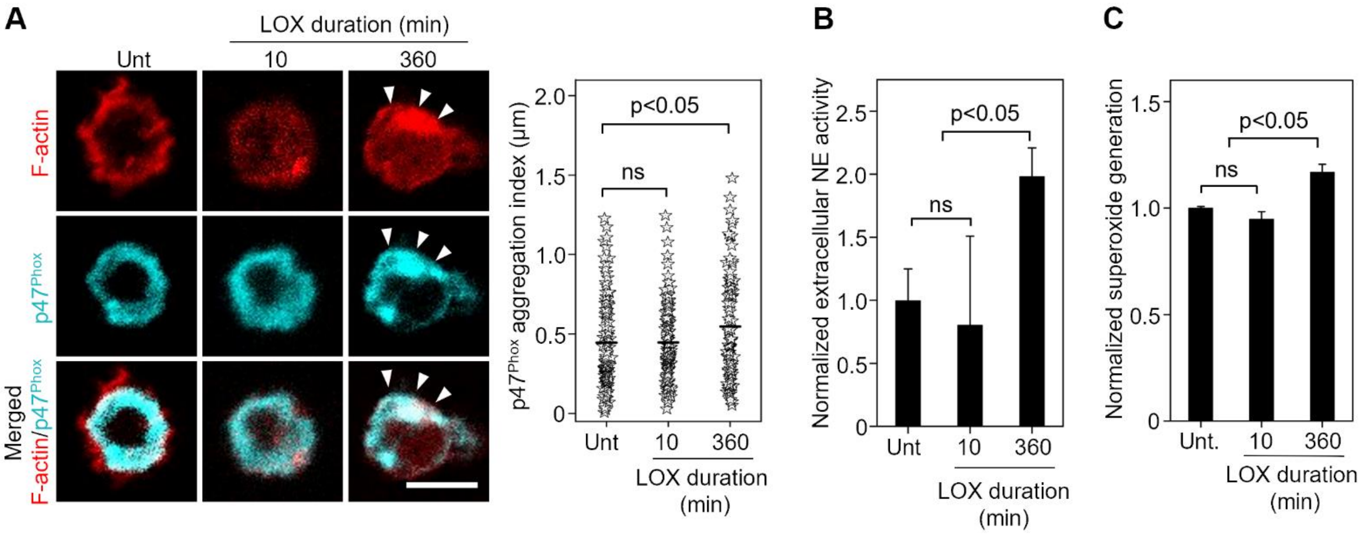


**Supplementary Fig 6: Temporal regulation of neutrophil activation and p47$^{Phox}$ aggregation by LOX. (A)** Untreated (Unt.) or LOX-treated dHL-60 cells were labelled with Phalloidin-594 (*red*) and anti-p47$^{Phox}$ (to visualize assembly of NADPH complex; *cyan*). Representative confocal images and subsequent quantitative analysis demonstrate that LOX-induced F-actin polarization is spatiotemporally associated with p47$^{Phox}$ aggregation (arrowheads) at 360 min (6h). **(B)** dHL-60 cells treated with 75 ng/mL recombinant LOX for 10 or 360 min were briefly stimulated with fMLP (10 nM) prior to supernatant collection. Fluorometric analysis of the supernatant/EnzChek™ elastase substrate mix revealed that LOX induces a significant increase in extracellular NE activity at 360 min (6h), but not 10 min. **(C)** dHL-60 cells treated with recombinant LOX (75 ng/mL; 6h) were resuspended in DHE solution prior to brief stimulation with fMLP (10 nM). Fluorometric analysis of the cell/DHE suspension revealed that LOX induces a significant increase in extracellular superoxide generation at 360 min (6h), but not 10 min. Column scatter plots indicate mean and distribution from ≥90 cells. Bar graphs indicate mean ± SEM. Scale bar, 5 μm.

**Supplementary Figure 7**

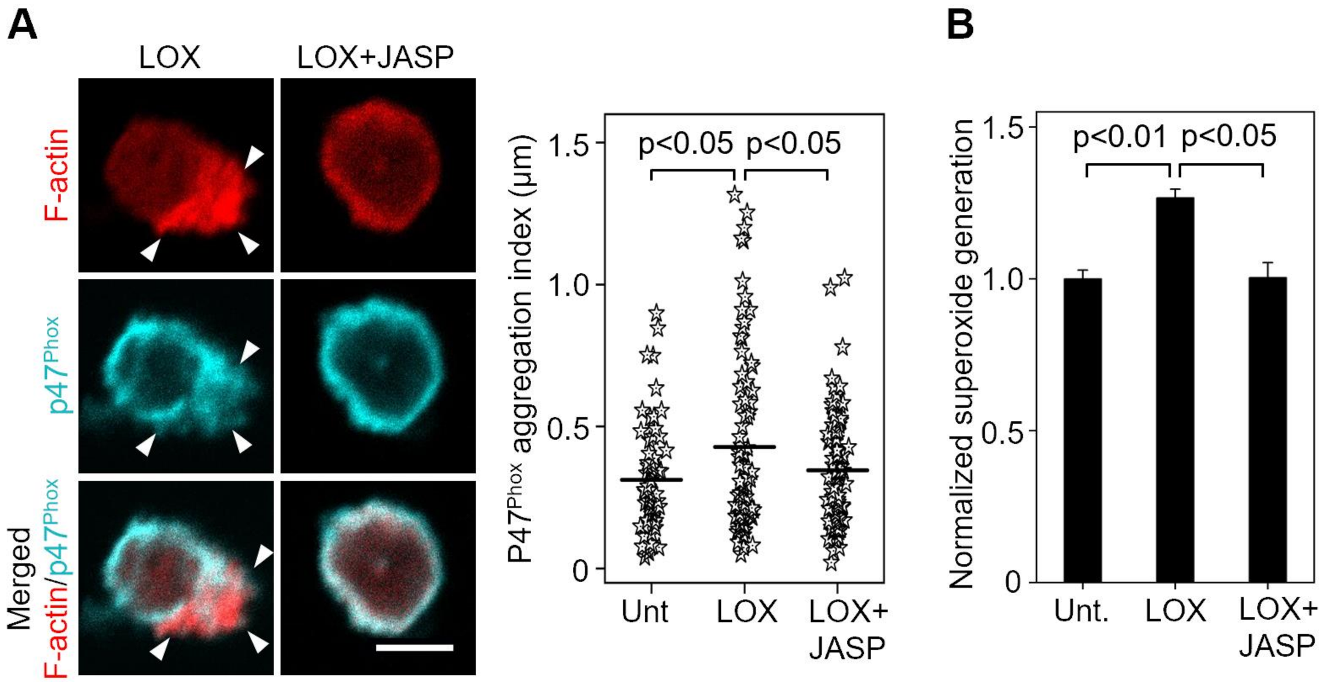


**Supplementary Fig. 7: Preventing transient actin depolymerization inhibits LOX-induced p47$^{Phox}$ aggregation and superoxide generation. (A)** dHL-60 cells treated with LOX (75 ng/mL) ± JASP (1 µM) for 6h were labelled with Phalloidin-594 (*red*) and anti-p47$^{Phox}$ (to visualize assembly of NADPH complex; *cyan*). Representative confocal images and subsequent quantitative analysis revealed that JASP treatment concurrently blocks LOX-induced F-actin polarization and p47$^{Phox}$ aggregation (arrowheads). Plots indicate mean and distribution from ≥65 cells. **(B)** Untreated (Unt.) or LOX-treated (75 ng/mL; 6h) dHL-60 cells were resuspended in DHE solution prior to brief stimulation with fMLP (10 nM). Fluorometric analysis of the cell/DHE suspension revealed that the LOX-induced increase in extracellular superoxide generation is completely blocked by JASP. Bar graph indicates mean ± SEM. Scale bar, 5 µm.

**Supplementary Figure 8**

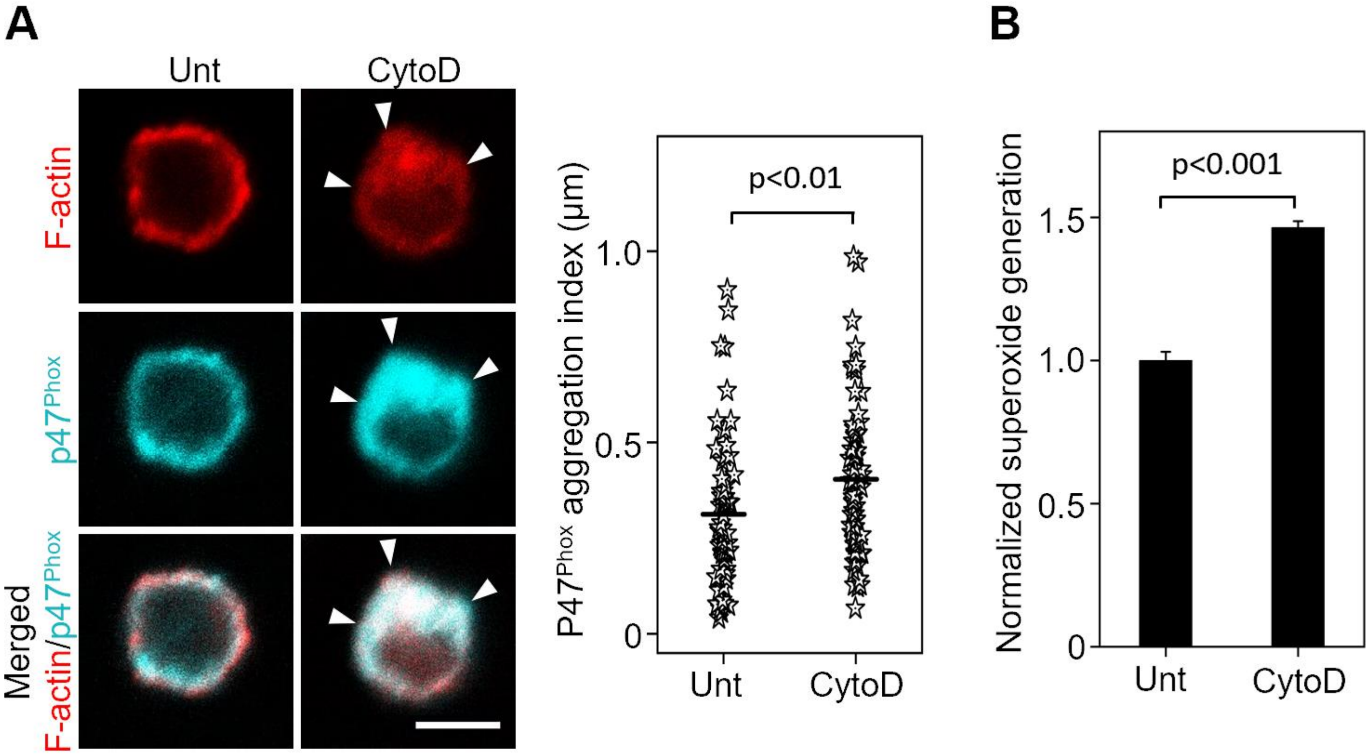


**Supplementary Fig 8: Transient actin depolymerization is sufficient to induce p47$^{Phox}$ aggregation and superoxide generation.** dHL-60 cells treated with CytoD (2.5 μM; 6h) were either (A) labelled with anti-p47$^{Phox}$ (to visualize assembly of NADPH complex; *cyan*) and Phalloidin-594 (to visualize F-actin; *red*), or (B) assessed for extracellular superoxide generation. (A) Representative confocal images and subsequent quantitative analysis revealed that CytoD treatment induces co-localized F-actin polarization and p47$^{Phox}$ aggregation (arrowheads) at 360 min (6h). Plot indicates mean and distribution from ≥65 cells. (B) Fluorometric analysis of the cell/DHE suspension revealed that CytoD treatment causes a significant increase in extracellular superoxide generation at 360 min (6h). Unt: Untreated. Bar graph indicates mean ± SEM. Scale bar, 5 μm.

**Supplementary Figure 9**

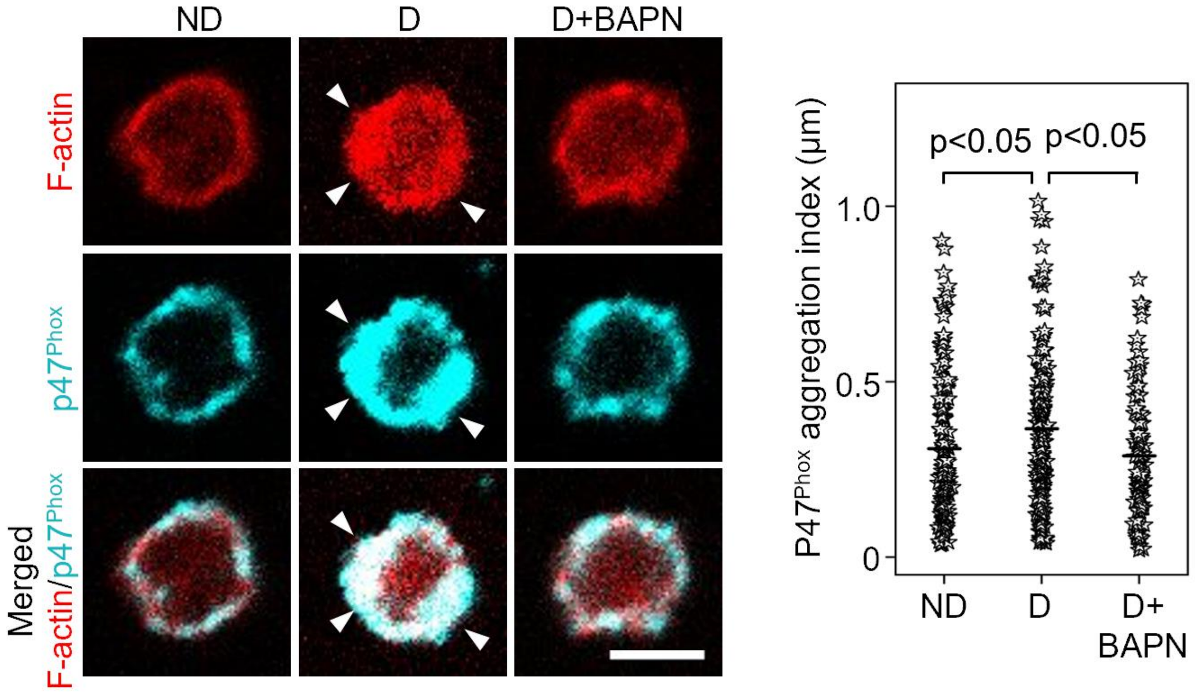


**Supplementary Fig 9: LOX inhibition blocks diabetes-induced p47$^{Phox}$ aggregation in neutrophils.** Mouse bone marrow neutrophils isolated from nondiabetic (ND), diabetic (D), or D mice treated with BAPN (D+BAPN; 3 mg/kg BW) were labelled with Phalloidin-594 (to visualize F-actin; *red*) and anti-p47$^{Phox}$ (to visualize assembly of NADPH complex; *cyan*). Representative confocal images and subsequent quantitative analysis revealed that LOX inhibition using BAPN blocks the diabetes-induced co-localized F-actin polarization and p47$^{Phox}$ aggregation (indicated by arrowheads). Plots indicate mean and distribution from ≥90 cells. Scale bar, 5 µm.
